# Flexibility of cell fates and functions across sex determination systems revealed by comparative single-cell analyses

**DOI:** 10.64898/2026.01.29.701242

**Authors:** Rafael D. Acemel, Boris Tezak, Vicky Wai Yee Chung, Alicia Hurtado, Scarlett Albertson, Ceri J. Weber, Blanche Capel, Darío G. Lupiáñez

## Abstract

Sex determination in vertebrates can be initiated by a wide range of genetic or environmental triggers. Yet, the degree to which gonadal cell types and genetic programs are conserved remains unresolved. Here we employed single-cell transcriptomics to characterize the temperature-dependent sex determination (TSD) program in gonads from the turtle species *Trachemys scripta.* Comparative analyses against species with genetic sex determination, like mouse (XY) and chicken (ZW), revealed a marked divergence in cell type repertoires and functions during vertebrate evolution. Unlike mammals, fetal Leydig cells are absent from the early gonads of *T. scripta*, where the supporting lineage expresses genes required for androgen synthesis. Evolutionary reconstructions show that this lineage derives from a *Pax2*-positive mesenchymal population, suggesting an ancestral condition in Archelosauria that differs from the primarily coelomic epithelium origin in the mammalian clade. Transcriptional dynamics and co-expression analyses revealed the recruitment of lineage-specific transcription factors, including *Twist1 or Runx1*, into the genetic programs of vertebrate clades. Our findings reveal extensive plasticity of the cellular and genetic mechanisms of vertebrate sex determination and suggest that this flexibility is a key feature of gonadal evolution.

## INTRODUCTION

Sex determination is the process by which an initially bipotential gonad commits to either the ovarian or testicular fate. This process is paramount for sexual reproduction but, despite this importance, sex determination mechanisms can evolve fast even within vertebrates ^1^. In animals with genetic sex determination (GSD), sex is determined by the presence or absence of a sex chromosome. That is the case for mammals that display an XY sex determination system (males are XY) ^2,3^ or birds which have a ZW sex determination system (females are ZW) ^4^. In addition to the different varieties of GSD, numerous vertebrate species determine sex through environmental cues that can include social interactions, food availability or pH. One of the most prevalent environmental mechanisms is temperature-dependent sex determination (TSD), which has been described in a wide range of animal species including turtles and alligators ^5^. Despite the remarkable variety of inputs that can initiate sex determination, the gonads of vertebrates share similarities at the morphological, physiological and gene expression levels. ^6^ Those observations led to the hypothesis that the cell types, developmental processes and genetic programs underlying sex determination are mostly conserved, even when they may respond to different initial triggers.^7^

Recent single-cell RNA-seq studies in model organisms, like mouse ^8,9^ or chicken ^10^, have contributed to finely delineate the cell types and lineages involved in sex determination. Three main cell lineages are distinguished: germ cells, supporting cells and interstitial cells. Supporting cells, which differentiate into either Sertoli in males or granulosa cells in females, are of special interest. They pioneer sex determination in all studied vertebrate species and instruct the rest of the gonad, including germ cells, towards the testicular or the ovarian fate ^11^. Importantly, the supporting cell lineages of mice and chickens have a different developmental origin. In mouse gonads, Sertoli and granulosa cells derive primarily from the coelomic epithelium ^8,9,12^, while in chicken and other birds they originate from mesenchymal cells that express the transcription factor (TF) Pax2 ^10,13^. Interestingly, a new cell type called supporting like cells (SLC) has been recently described in mammals ^14^. These cells are similar transcriptomically to supporting cells, but express the TF PAX8. SLCs mostly contribute to the formation of the rete testis and rete ovarii, but they also constitute a minor source of Sertoli and granulosa cells. Of relevance are also the fetal Leydig cells of the mammalian testis, derived from the interstitial lineage ^15^, which are responsible for the production of androgens from cholesterol during prenatal stages ^16^. Estrogens, produced through the aromatase-mediated conversion of androgens^17^, have been also reported to be synthesized in the prenatal ovaries of some mammalian species, such as human or rabbits, although the specific cell type responsible for this activity remains uncertain. While sex steroids are dispensable for sex determination in mammals ^18^, they play a more prominent role modulating gonadal fate in non-mammalian vertebrate species, with their manipulation often leading to gonadal sex reversal. That is the case in chicken, where androgens are produced by fetal Leydig cells in testes and theca cells in ovaries, but converted into estrogens exclusively in the ovaries by granulosa cells. Importantly, Leydig cells and theca cells in the chicken appear to derive from supporting cells and not from the interstitial lineage, as described for mammals ^10^. Taken together, these findings indicate divergence in the origin of supporting and steroidogenic gonadal cell types across vertebrate evolution.

To extend our understanding of the evolution of the gonad to vertebrates that lack sex chromosomes and instead rely on TSD systems, we directed our attention to the red-eared slider turtle *Trachemys scripta*. *T. scripta* displays a warm-female/cool-male TSD system, so constant warm (31°C) incubations give rise to female offspring while cool (26°C) incubations promote exclusively male offspring ^19^. Temperature and sex are linked via the phosphorylation of Stat3 in response to the influx of calcium at 31°C. pSTAT3 then promotes ovarian differentiation through the repression of *Kdm6b* and its target, *Dmrt1* ^20^. Additionally, sex hormones can play a pivotal role, as demonstrated by the sex reversal phenotypes obtained upon pharmacological or genetic manipulation of androgen and estrogen levels ^21,22^. Here, to clarify the evolution of the cell types and genetic programs underlying gonadal sex determination and differentiation, we employed single-cell RNA-seq to investigate cell lineages and their origins during TSD in *T. scripta.* Crucially, we put our results in the context of available datasets from vertebrates with GSD (chicken ^10^ and mouse ^8,9^). We found that, although coarse cell lineages are conserved among all vertebrates studied, striking differences were evident at the level of cell types and origins, cell differentiation mechanisms and genetic programs.

## RESULTS

### Transcriptional dynamics during TSD in single cells

To shed light on the cellular and molecular mechanisms behind sex determination in response to temperature, we carried out single-cell RNA-seq experiments in gonads from *T. scripta* embryos around the thermosensitive period. Embryos were incubated at either the male-promoting temperature (MPT) of 26 °C or the female-promoting temperature (FPT) of 31°C (Fig. 1a). Gonadal samples were collected at Greenbaum Stg.15, Stg.18 and Stg.23 which corresponds to undifferentiated, sex-determining and differentiated gonads (testes and ovaries respectively). Altogether, we sequenced a total of 25,748 cells from six different conditions. Single-cell transcriptomes from the different experiments were integrated for dimensionality reduction and clustering, obtaining 26 different clusters (see methods, Extended Data Fig. 1a). Consistent with *a priori* expectations, we found more similar cell compositions among Stg.15 gonadal samples raised at either MPT or FPT (prior to the sex determination window). In contrast, gonads from the latest stage Stg. 23 displayed clusters of cells that were clearly sex-specific (Fig. 1b, Extended Data Fig. 1b).

**Figure 1.**
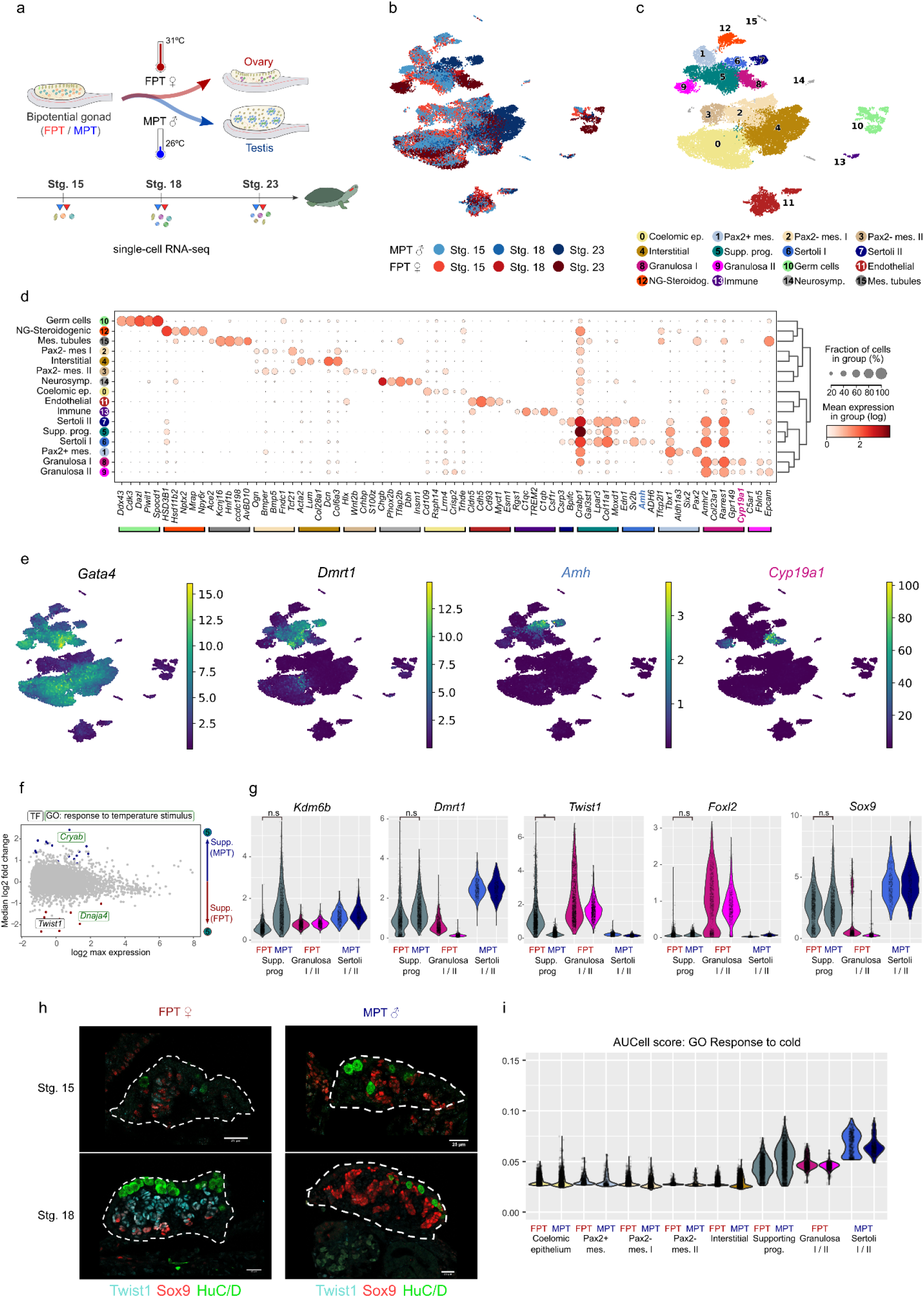
Temperature-dependent sex determination at the single cell scale. a) Diagram of bipotential gonad differentiation in either testis or ovaries in response to cold MPT or warm FPT respectively. Cells from critical stages before, during and after the sex determination were collected from single-cell RNA-seq. b) UMAP projection after the integration of the six different samples: 3 stages at FPT and 3 stages of MPT. Lighter colors represent earlier stages. c) Same UMAP projection colored by the different cell types identified after Leiden clustering. d) Dotplot displaying the expression of the top 3 non-redundant markers per cell type. The size of the dot represents the percentage of the cells expressing the gene in each cell type and the color the mean expression. e) SCVI normalized expression of the genes *Gata4, Dmrt1, Amh* and *Cyp19a1* projected on the UMAP. f) MA plot from the differential expression analysis between MPT and FPT in the cell type Supporting progenitors. The x-axis represents the logarithm of the maximum mean expression and the y-axis the fold change. Negative values of fold changes correspond to genes more expressed in FPT. Genes upregulated in FPT are highlighted in red and genes upregulated in MPT in blue. Transcription factors (TFs) and members of the GO term *response to temperature stimulus* are squared black and green respectively g) SCVI normalized expression of *Kdm6b, Dmrt1, Twist1, Sox9* and *Foxl2* in cell types of the supporting lineage. h) Representative immunofluorescence images of gonadal cross sections from St.15 and St.18 incubated at FPT and MPT. Twist1 is labeled in cyan, Sox9 in red, and HuC/D (labelling germ cells) in green. The gonad is outlined with white dashed line (scalebar, 25 μm). i) Combined AUCell scores of the gene set composed by the genes belonging to the GO term *response to cold* in each cell type at either FPT or MPT.

To assign identity to each gonadal cell cluster we performed differential expression analysis to obtain marker genes (Extended Data Fig. 2), focusing on transcription factors (TF). Using a combination of GSEA GO enrichment analysis (Extended Data Fig. 3) and available literature we identified 16 different cell types with clearly distinct transcriptional signatures (Fig. 1c-d, Extended Data Fig. 2). Germ cells, studied extensively in a previous publication ^23^, were recognized by the expression of known markers like *Dazl* or *Piwi1* and the enrichment of GO terms like *meiotic cell cycle*. Cell types belonging to the somatic portion of the gonad were identified using the known marker *Gata4* (Fig. 1e). Among the *Gata4+* cells we identified (i) cells from the coelomic epithelium expressing epithelial markers like *Cd109* and high levels of the TFs *Lhx9* and *Emx2,* (ii) interstitial cells expressing collagen and extracellular-matrix related genes (i.e. *Col6a3* and *Dcn*) and (iii) cells from the supporting lineage. Supporting progenitor cells from early stages are distinguished by the expression of *Dmrt1* and *Sox9* (Fig. 1e). These TFs continue to be expressed in the two clusters of Sertoli cells, characterized also by the expression of the *Amh* gene encoding for the anti-mullerian hormone (Fig. 1e). In contrast, *Dmrt1* and *Sox9* are downregulated in the two distinct clusters of granulosa cells which express the aromatase encoding gene *Cyp19a1* (Fig. 1e) and the TF *Foxl2*. We also identified *Gata4*-negative (*Gata4-*) cell types like *Cldn5/Sox18*+ endothelial cells (enriched in angiogenesis GO terms) and mesenchymal cell populations that were either *Pax2/Tbx1*+ (*Pax2+* mes.) or *Tcf21*+ (*Pax2-* mes. I and II, enriched in extracellular matrix remodelling GO terms). Importantly, *Gata4*-mesenchymal cell clusters were only observed at early stages of both MPT and FPT gonads. Finally, additional clusters of extragonadal cell types were also distinguished. Those include *Mrap/Hsd11b2*+ non-gonadal steroidogenic cells (expressing genes related with melanocortin and cortisol signalling), *Phox2b/Tfap2b*+ sympathetic peripheral neurons (GO term *neurotransmitter transport*), *Hnf1b/Kcnj16+* mesonephric tubule cells (enriched in GO terms related to ion transport) and *C1qb/C1qb+* immune cells (immune system related GO terms). We further validated relevant signatures within *T. scripta* gonads through immunofluorescence, confirming the spatial distribution of the somatic *Gata4*+ populations, germ cells and the supporting lineage (Fig. 1h; Extended Data Fig. 4).

After assigning cell type identities, we investigated early differences between FPT and MPT gonads in the supporting progenitors to identify uncharacterized TSD determinants in turtles. We performed differential expression analysis and identified only 9 and 16 genes upregulated in FPT and MPT supporting cells respectively (Fig. 1f). Previous works proposed that the first sexually dimorphic event at the transcriptional level in *T. scripta* is the downregulation of *Kdm6b* and *Dmrt1* in FPT in response to phosphorylated Stat3 (pSTAT3)^20^. Consistently, we found an MPT biased expression of *Kdm6b* and *Dmrt1* in supporting progenitors, although differences were not statistically significant (Fig. 1g). Next we manually inspected the 25 differentially expressed genes (DEG) across all clusters and pinpointed those genes whose differences in expression were restricted to the supporting lineage (Extended Data Fig. 5). We found 2 FPT biased genes (*Twist1, Dnaja4*) and 5 MPT biased genes (*Atp6ap2*, *Cryab*, *Adm2*, *Susd1* and *Gfra3*). We specifically focused on *Twist1* as it is the only transcription factor that was significantly upregulated in our single-cell analysis (Fig. 1g) and it has not been linked to sex determination. Furthermore, *Twist1* expression in female supporting cells precedes *Foxl2* expression and Sox9 downregulation (Fig. 1g), suggesting that it operates early. We characterized the expression pattern of Twist1 through immunofluorescence (Fig. 1h, Extended Data Fig. 6a) and confirmed its FPT-biased expression. In the supporting lineage of FPT gonads, Twist1 expression starts from Stg.15 and peaks at Stg.18, gradually decreasing expression after Stg.19 when the gonad starts to display clear patterns of ovarian differentiation (Extended Data Fig. 6b). In contrast, Twist1 was not detected via immunofluorescence in MPT gonads from Stg.15-23. Co-immunofluorescence with the testicular marker Sox9 revealed that the expression of Sox9 precedes Twist1 expression in the supporting lineage of FPT gonads at early stages. Yet, both markers colocalize extensively through Stgs.16-17 and resolve to Twist+/Sox9-cells at FPT Stgs.18-19. These dynamic expression patterns are compatible with an antagonistic relationship between Sox9 and Twist1, as it has been reported during chondrogenesis in mouse ^24^. Next, we explored the conservation of *Twist1* expression in supporting cells in other vertebrate species by evaluating its expression in publicly available datasets (Extended Data Fig. 6c-d). We found that *Twist1* was not expressed in supporting cells in chicken or mouse (Extended Data Fig. 6e-f), suggesting that a putative role in sex determination could be restricted to turtles.

To identify sexually dimorphic biological processes, we used Gene Set Enrichment Analysis (GSEA) in supporting cells at either FPT or MPT. This analysis revealed 106 GO terms upregulated between gonads incubated at distinct temperatures (Extended Data Fig. 7). At FPT, we found 22 GO terms enriched related to core cellular processes like synthesis of DNA and protein and cellular respiration. In contrast, the 84 terms enriched at MPT revealed the activation of processes related to morphogenesis and development (i.e. *cell morphogenesis, cell migration, cell junction organization* or *negative chemotaxis*) and with the physiology of more mature Sertoli cells (i.e. *response to hormone, lipid transport, regulated exocytosis*). Suggestively, GO terms like *MAPK cascade* (involved in male sex determination in mice ^25^), *phosphatidylinositol metabolism* and *response to endoplasmic reticulum stress* (both tightly linked with Ca^2+^ signalling) were also upregulated in MPT supporting cells (Extended Data Fig. 7). The latter could be activated in response to defects in protein folding following cold incubation temperatures. Indeed, among the DEGs we found two genes (*Cryab* and *Dnaja4*) related to *protein folding* and *response to temperature* which prompted us to investigate the molecular response to temperature further. We found individual genes involved in processes like *response to temperature stimulus*, *response to heat* and *response to cold* that were expressed across all gonadal cell types (Extended Data Fig. 8a), but were most highly represented in the supporting lineage (Extended Data Fig. 8b). In particular, the combined expression of *response to cold* genes (calculated with AUCell ^26^) was higher in MPT supporting lineage cells in comparison with FPT cells or MPT cells from other cell types (Fig. 1i). This suggests that cells of the supporting lineage might be particularly sensitive to temperature changes. To investigate this aspect further, we explored the expression of the temperature dependent Trpv Ca^2+^ channels, implicated in TSD in alligators ^27^. We found that *Trpv4* and *Trpv5-6* were expressed in supporting cells (Extended Data Fig. 8c), suggesting a conserved role as temperature sensing channels in TSD.

### Evolution of the genetic programs across vertebrate gonadal cell types

Sex determination mechanisms diverge among vertebrate clades (i.e. TSD in *T. scripta*, XY GSD in mammals and ZW GSD in birds). Yet, our initial comparisons of single-cell data from gonads of turtle, mouse and chicken suggested homologies across most gonadal cell types, as denoted by the expression of common marker genes (Fig. 1c, Extended Data Fig. 6c-d). Although it was possible to infer the overall conservation of the main critical lineages, germ cells, supporting cells and interstitial cells, notable exceptions were mouse fetal Leydig cells and theca cells of chicken ovaries, which had no obvious counterparts in turtle gonads.

To investigate the degree of conservation of the genetic programs that define the identity of critical gonadal cell types we used Weighted Correlation Network Analysis (WGCNA) ^28,29^. First, we focused on the comparison between gonads of turtles and chickens, as these species are closer phylogenetically. We obtained 10 WGCNA coexpression modules in chickens ^10^ (Fig. 2a), together with their consensus expression profiles known as their module eigengenes (ME). This metric helped us to recognize that 9 out of the 10 modules were associated with particular cell-types (Extended Data Fig. 9a) and were named accordingly (Fig. 2a). To further characterize WGCNA modules, we explored the genes associated with them by ranking them according to the similarity of their expression profile to the ME of each module, a metric referred to as the gene ME connectivity (kME). Genes with high kME values are more associated with a particular module and were thus considered as hub genes. This analysis facilitated the identification of the module without a clear association with a cell type, as a cell cycle module (Extended Data Fig. 9b). In addition, the identification of hub genes confirmed the presence of known regulators of gonad development within the different WGCNA modules. For instance, *Foxl2/Cyp19a1/Sall1* appeared as hub genes of the *granulosa* module, *Amh/Sox9* in *Sertoli,* and the androgen producing enzymes *Star/Cyp11a1/Cyp17a1* in the *Theca* module (Extended Data Fig. 9b). Next, we explored the degree of conservation of chicken WGCNA modules in turtles. For this purpose, we used 1-to-1 orthologs to recalculate ME in our single-cell datasets from turtle gonads. We discovered that 5 out of the 9 modules (*Supporting lineage, Sertoli, granulosa, Interstitial prog.* and *Interstitial)* displayed higher ME values in the expected putatively homologous cell type (Fig. 2b), suggesting at least a partial conservation of cell types and genetic programs across species. Of the 4 remaining modules, 3 were related with the coelomic epithelium and the ovarian cortex of chicken which are highly specialized (*Coelomic epithelium I* and *II* and *Ovarian cortex*). The remaining module was the *Theca,* which was not associated with a particular cell type as observed in chicken. In contrast, this module displayed higher ME scores across all the cell types of the turtle supporting lineage, suggesting a distribution of the steroidogenic role among several cell types.

**Figure 2.**
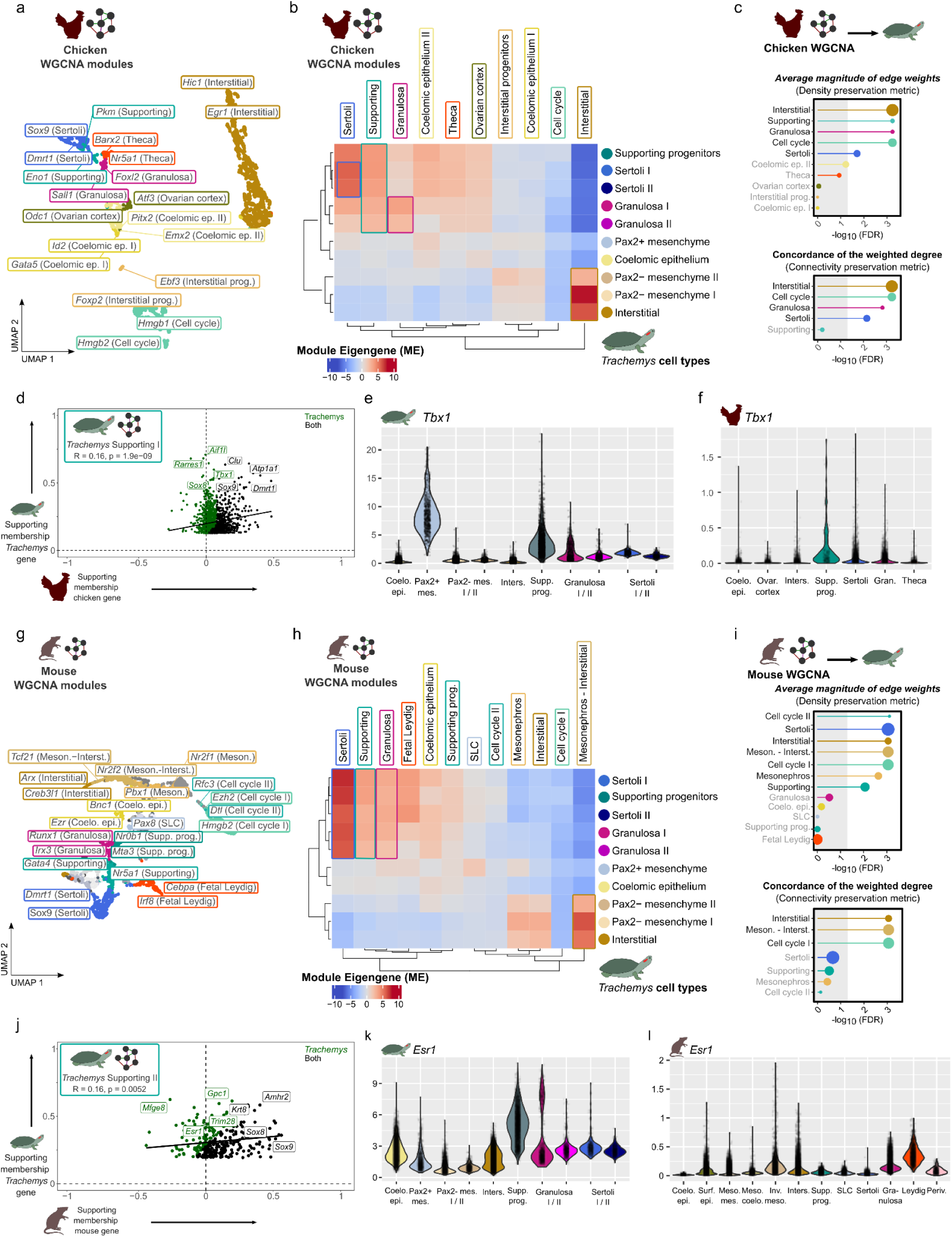
The evolution of the genetic programs of gonadal cell types in vertebrates. a) UMAP projection in which each dot represents a chicken gene in a WGCNA module. Coexpressed genes are grouped together. The name of the top TFs per module are shown. b) Heatmap showing Module Eigengene (ME) scores of chicken WGCNA modules in each of the *T. scripta* cell types. High ME scores imply that the genes of the chicken WGCNA module are active in the corresponding cell type of *T. scripta*. Rectangles highlight chicken WGCNA modules that are active in putatively homologous cell types in both chicken and *T. scripta*. c) Lollipop plot displaying network conservation FDR values between chicken and *T. scripta* of the different WGCNA modules calculated with NetRep. On the x-axis is represented the negative logarithm of the FDR of a permutation test assessing whether the module has a higher conservation metric than modules generated by chance. Density conservation metrics (top) assess whether the genes that are coexpressed in the WGCNA module in chicken are also coexpressed in *T. scripta.* Connectivity preservation metrics (bottom) assess whether the hierarchy of the module is conserved, e.g the genes more central to the module in chicken are also central in *T. scripta*. The size of the lollipop circle is proportional to the number of genes in the module. d) Scatterplot showing the membership (kME) of the genes belonging to the WGNCA module *Supporting I* calculated in *T.scripta* both in *T.scripta* (y-axis) and chicken (x-axis). Most of the genes in the module have kME scores that are greater than 0 in chicken indicating that the module is conserved (labeled black, i.e. *Dmrt1*, *Sox9*). Genes that are associated in *T. scripta* but not in chicken (KME differences above the third quartile plus 1.5 times the interquartile range) are labeled green (i.e. *Tbx1*). e-f) SCVI normalized expression of *Tbx1* in *T.scripta* and chicken cell types respectively. g) As in a), UMAP projection of the genes grouped by WGCNA module calculated in mouse. h) As in b) heatmap showing ME scores of mouse WGCNA modules in *T. scripta* cell types. i) As in c) lollipop plot showing NetRep network conservation metrics of mouse WGCNA modules in *T. scripta*. j) As in d) for the genes of the *T. scripta* module *Supporting II* in *T.scripta* (y-axis) and mouse (x-axis). k-l) SCVI normalized expression of *Esr1* in *T. scripta* and mouse cell types respectively.

To further investigate the cross-species conservation of WGCNA modules, we obtained conservation metrics using NetRep ^30^. These metrics fall into two main categories: density and connectivity. In this context, density metrics evaluate whether the genes grouped into a WGCNA module in one species remain coexpressed in the second species. In contrast, connectivity metrics evaluate whether the relationships between the genes in the module are maintained (i.e. their relative importance). It is worth noting that the conservation at the connectivity level of a module that is not conserved at the density level is difficult to interpret biologically, and therefore we focused on density metrics first. Here, 5 out of the 10 WGCNA modules calculated in chicken were found to be conserved in turtles at the density level (*Supporting, Sertoli, granulosa, Interstitial* and *Cell cycle*) and 4 of them were also conserved at the connectivity level (excluding *Supporting,* Fig. 2c, Extended Data Fig. 10). We then performed the converse analysis and identified 17 WGCNA modules in turtles. A total of 9 modules could be assigned to one or several turtle cell types, while the remaining ones displayed a pervasive distribution across cell types, including a cell cycle module (Extended Data Fig. 11). 6 out of the 9 cell-type specific modules displayed higher ME values in the homologous chicken cell types, including the *Coelomic epithelium* module which corresponded to the Coelomic epithelium and Ovarian cortex clusters (Extended Data Fig. 12a). Seemingly inconsistent with these results, NetRep conservation analysis only identified three WGCNA modules of turtles conserved in chicken at the density level (*Supporting I*, *Mesonephros-Interstitial* and *Cell cycle*) and the module *Supporting I* was not conserved at the connectivity level (Extended Data Fig. 12b). Unexpectedly, the modules *Sertoli* and *granulosa* calculated in chicken were conserved in turtles and not vice versa. We hypothesized that core genetic programs identified in the Sertoli and granulosa cells of chicken are conserved also in *T. scripta*. However, additional genes were incorporated independently in the turtle supporting cell lineage modules.

In order to pinpoint genes that were incorporated specifically into *T. scripta* genetic programs, we focused on genes that were highly associated to a turtle module but lost this association in chicken. This analysis revealed several examples of turtle specific genes like the TF *Tbx1,* which is highly associated with the *Supporting I* module in turtles but not in chicken (Fig. 2d). Accordingly, *Tbx1* was expressed specifically in supporting progenitor cells and *Pax2*+ mes. cells in turtles and lacks this cell type specificity in chicken (Fig. 2e-f). Likewise, the TF *Nkx3-1* and the signalling molecule *Wnt16* were exclusively associated with the *T. scripta* modules *Sertoli* and *granulosa* respectively (Extended Data Fig. 12c-h). Hence, cross-species WGCNA analysis not only allowed us to identify which genetic circuits are conserved, but to single out species-specific acquisitions associated with cell type evolution.

Next, we wondered if the moderate conservation of modules and cell types between turtles and birds could be also expanded to mammals. We performed analogous analyses in mouse gonads and found 32 WGCNA modules, 10 of them being associated with cell types (Fig. 2g, Extended Data Fig. 13). Notably, when we assessed the expression of these modules of genes in turtles we could recognize coarse cell lineages (Fig. 2h, Extended Data Fig. 14a). For instance, ME values for the mouse modules Sertoli and granulosa were not directly associated with the corresponding cell types in turtles, but rather peaked equally in all cell clusters belonging to the supporting lineage. Similarly, the mouse modules *Mesonephros, Interstitial* and *Mesonephros-interstitial* peaked in both turtle populations. Taken together, these data suggest the presence of homologous cell lineages and genetic circuits operating in both *T. scripta* and mammals. Nevertheless, as expected, those lineages and circuits are more divergent than in the comparison with chicken. Indeed, we interrogated the conservation of the coexpression modules using NetRep and found that 5 out of the 10 modules (*Supporting, Sertoli*, *Interstitial*, *Mesonephros*, *Mesonephros-interstitial*) were conserved at the density level, however, only the last one (*Mesonephros-interstitial*) was conserved at the connectivity level (Fig. 2i, Extended Data Fig. 14b). Among the modules with lower conservation scores we found both the *Fetal Leydig* cell, with no clear counterpart in turtles, but also the *granulosa* module. We analyzed the source of the lack of conservation of the module *granulosa* in turtles (Extended Data Fig. 14c). Among other genes, the hub TFs *Runx1* and *Irx3,* were absent or lowly expressed in *T. scripta* granulosa cells during the sex determination window (Extended Data Fig. 14d-g). This finding was surprising because the TF *Runx1* has a well studied, conserved role in the specification of granulosa cells, from mammals to teleost fishes ^31^. We took advantage of a well defined list of Runx1 targets in mouse ovaries to further explore the potential dismantling of the Runx1-driven genetic program in *T. scripta.* To this end we interrogated the expression of RUNX1 targets in mouse and turtles embryonic gonads (Extended Data Fig. 15a-b). In mice, the combined expression of RUNX1 targets mirrors that of the transcription factor: active in granulosa cells and supporting progenitors and absent from Sertoli cells. In contrast, in *T. scripta* gonads where Runx1 is not detected, Runx1 targets are expressed broadly and the female specific expression in granulosa cells is lost. Additionally, we could identify a set of 12 targets (i.e. *Ryr2, Mmp15, Mettl24*) that were clearly associated with the module *granulosa* in mice but not in turtles (Extended Data Fig. 15c). These analyses suggest that *Runx1* may have lost its role as an ovarian differentiation factor in turtles.

Finally, we performed the converse analysis, and recalculated WGNA modules in turtles to assess their conservation in mice (Extended Data Fig. 16a-b). We found that only the module *Mesonephros-Interstial* was conserved both at the density and connectivity level and the *Supporting I* only at the density level (driven by common hub genes such as *Dmrt1*, *Sox9* and *Nr5a1,* Extended Data Fig. 16c). The divergence between modules of the supporting lineage between mouse and turtles prompted us again to identify turtle-specific genetic modules involved in sex determination. In the case of Sertoli cells, we found again the turtle-specific association of the transcription factor *Nkx3-1* with the *Sertoli* module (Extended Data Fig. 16d-e). In the more divergent module *granulosa* (Extended Data Fig. 16f), genes such as *Ar* and *Cyp19a1* were also uniquely associated in turtles (Extended Data Fig. 16 g-j). Finally, *Esr1* showed a turtle-specific association with the module *Supporting II* (Fig. 2j-l). The central role of genes related to the synthesis (*Cyp19a1*) and response (*Ar*, *Esr1*) of androgens and estrogens in turtles reflect the importance of steroid hormones for sex differentiation in non mammalian vertebrates. Taken together, WGCNA results indicate that the genetic programs responsible for the specification of critical cell lineages in sex determination across vertebrates are sufficiently similar to establish cell type homologies. However, they have been heavily reconfigured in the different lineages independently, including the turnover of critical transcription factors. Illustrative examples of this turnover are the incorporation of Tbx1 or Nkx3.1 and the loss of importance of Runx1 or Irx3 in *T. scripta* gonads.

### Supporting cells mediate fetal steroidogenesis in turtles

Our WGCNA analyses revealed a lack of conservation of the steroidogenic modules *Theca* of chicken and *Fetal Leydig* of mouse, that were not assigned to any specific cell type of turtles (Fig. 2c,i). This result is remarkable, given that steroid hormones play during sex determination in turtles. To identify which cell populations are responsible for steroidogenesis in this species, we scored the expression of the genes of the modules *Theca* and *Fetal Leydig* in turtle cells. The ME scores of both modules indicated a moderate enrichment in clusters derived from the supporting lineage (e.g. Supporting progenitors, Sertoli and granulosa cells, Fig. 3a-b). For instance *Star*, which encodes the first enzyme required for androgen synthesis, was expressed specifically in the clusters Supporting progenitors, granulosa and Sertoli (Fig. 3c). Similarly, the genes that encode the enzymes that convert cholesterol into androgens (e.g. *Cyp11a1*, *Hsd3b* and *Cyp17a1*) were also predominantly expressed in cells of the supporting lineage (Fig. 3d-f). The aromatase gene *Cyp19a1*, responsible for the conversion of androgens into estrogens, was expressed exclusively in female granulosa cells (Fig. 3g). We employed immunofluorescence to validate the coexpression of the steroidogenic enzyme Cyp11a1 with the somatic and supporting cell markers Gata4 and Sox9 respectively in FPT and MPT gonads (Fig. 3h and Extended Data Fig. 17). In contrast to mammals, Cyp11a1 staining was found in supporting cells at both MPT and FPT within the primitive sex cords, denoted by its colocalization with Sox9, but not in the gonadal interstitium. Importantly we do not observe this colocalization outside the gonad, where Cyp11a1+ cells do not express Sox9. From an evolutionary perspective (Fig. 3i), androgens are produced in mammalian species by interstitial fetal Leydig cells and exclusively in males. In chicken, there are also cell types specialized in the synthesis of androgens, termed fetal theca in females and fetal Leydig in males ^10^. Yet, in contrast to mammals, both populations are derived from the supporting lineage (pre-Sertoli cells in males and pregranulosa cells in the female ^10^). In turtles the situation is more similar to chicken, since steroidogenic enzymes are also produced by supporting lineage cells. However, *T. scripta* seems to lack a specialized population undertaking androgen production. Rather, genes encoding steroid enzymes are expressed broadly in both undifferentiated supporting progenitor cells and more mature Sertoli and granulosa cells. These results show that analogous functions, in this case steroidogenesis, can be performed by different gonadal lineages across vertebrate clades.

**Figure 3.**
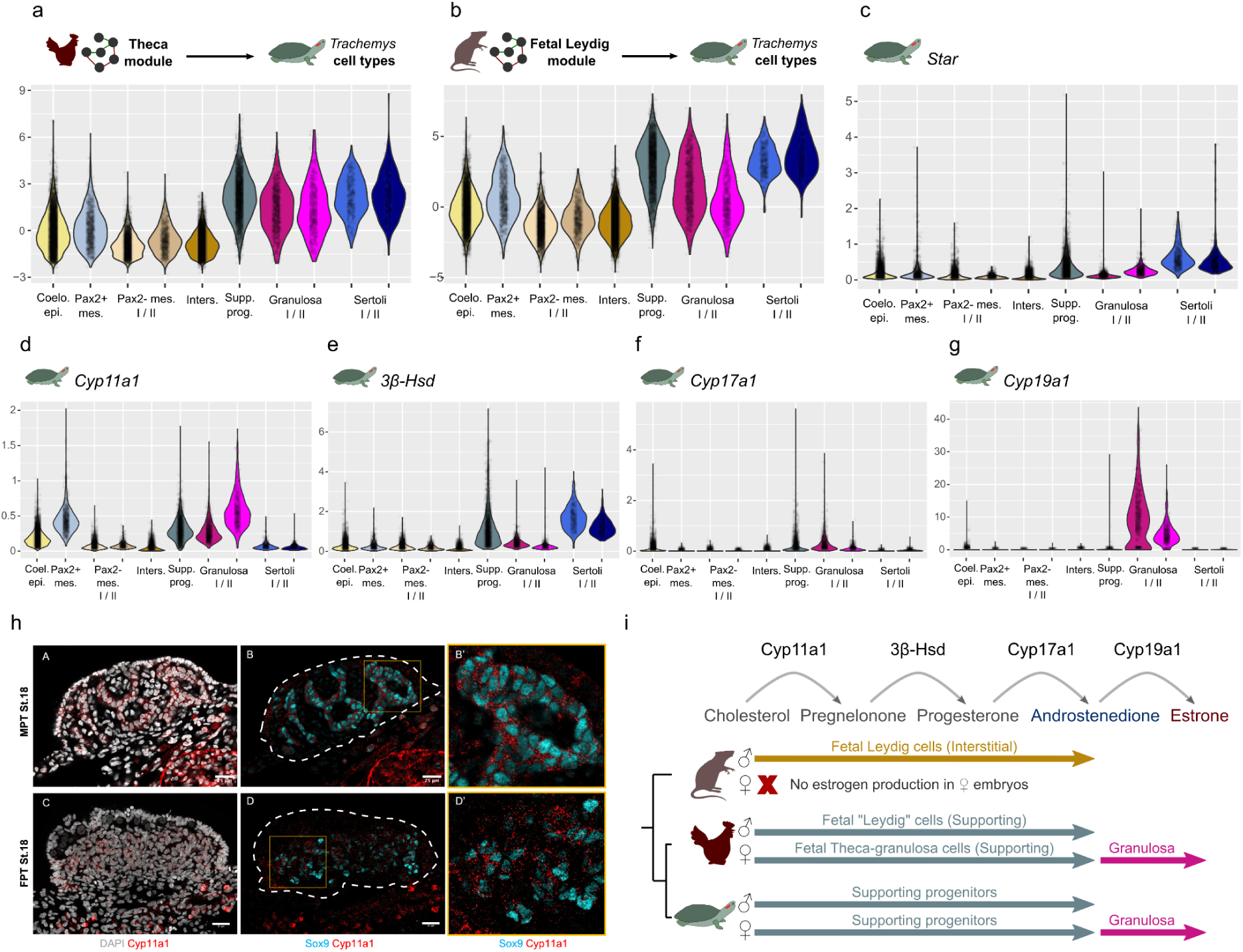
Steroidogenesis in *T. scripta* gonads. a-b) Module Eigengene (ME) scores of the WGCNA modules *Theca* and *Fetal Leydig* of chicken and mouse respectively in *T. scripta* cell types. c-g) SCVI normalized expression of genes encoding fundamental enzymes involved in the synthesis of androgens and estrogens. h) Representative immunofluorescence images of MPT and FPT gonadal cross sections at St.18. DAPI labels the nuclei in grey, Sox9 labels the supporting cells in cyan, and Cyp11a1 labels steroidogenic cells in red. Note that Cyp11a1 labels the cytoplasm of Sox9+ cells at both temperatures (scalebar, 25 μm) and that Cyp11a1+/Sox9-cells are located outside the gonad (delineated with a white dashed line). The yellow box depicts the region magnified on the right break-out panels. i) Diagram summarizing the process of sex hormone synthesis in different vertebrates with the cell type in charge of its production.

### A mesenchymal Pax2+ origin of the supporting lineage in Archelosauria

As described above, the supporting lineage in turtles is characterized by the expression of lineage-specific genes that have been extensively characterized in mammals, such as *Sox9* for Sertoli cells or *Foxl2* for granulosa cells. Yet, turtle supporting cells are also steroidogenic, denoting functional differences between homologous cell types across major vertebrate clades. Notably, the supporting lineage in chickens, which also has steroidogenic potential, originates from mesenchymal cells that express the transcription factor Pax2. ^10,13^ This contrasts with the developmental origin described in mice for this lineage. In this species, the supporting lineage mostly derives from the coelomic epithelium ^8,9,12^, with a minor contribution from SLC progenitors ^15,32^. Therefore, we explored whether the origin of the supporting cells in turtles is more similar to the condition described in mammals or birds.

Based on our clustering analyses at early stages, we could identify both potential sources for the supporting lineage: cells from the coelomic epithelium (*Lhx9*/*Emx2*+) and *Pax2*+ mesenchymal cells (Fig. 1c). While turtle *Pax2*+ mesenchymal cells also transcribe *Wnt4* and *Osr1* as in chicken (Extended Data Fig. 18a-b), they also display low levels of *Dmrt1*, thus denoting a certain degree of transcriptional divergence. To elucidate the origin of supporting cells in turtles we performed trajectory analysis in MPT and FPT cells separately (Fig. 4a-b). We used a modified version of the PAGA algorithm that takes into account the stage information to infer developmental trajectories (see Methods). Using the PAGA graphs, we calculated the shortest distances from each early-stage cell cluster to each late-stage cluster. As a control, we applied our modified PAGA algorithm to available mouse and chicken datasets and could confirm the previously validated origin of Sertoli and granulosa cells (Pax2+ cells in chicken and coelomic epithelium cells in mouse, Extended Data Figures 19-20). In turtle FPT data, we observed that granulosa cells were closer to *Pax2*+ cells than to coelomic epithelium cells in PAGA graph distances (Fig. 4a); whereas, the coelomic epithelium cells were more related to the ovarian cortex epithelium (Fig. 4a, Extended Data Fig. 18c,e). Similarly, in MPT samples, we observe that Sertoli cells were closer to *Pax2*+ than to coelomic epithelium cells (Fig. 4b, Extended Data Fig. 18d,f). In fact, coelomic epithelium cells were closer to terminal clusters of interstitial cells than to Sertoli cells. Of note, *Pax2*-mesenchymal cells expressing the transcription factor *Tcf21* were predicted to be a main source of terminal Interstitial cells both in MPT and FPT (Extended Data Fig. 18c-f). To further investigate the *Pax2*+ mesenchymal origin of supporting cells in turtles, we performed immunofluorescence against Pax2 and Sox9 in early MPT and FPT gonads (Stgs.15 and 18). As described in birds, we identified the Pax2+ population adjacent to the mesonephros (Fig. 4c). Importantly, we identified individual cells coexpressing Pax2 and Sox9 at the mesonephric side of the sex cords (Fig. 4c), consistent with bioinformatic predictions that Sox9+ supporting cells in the sex cords derive from Pax2+ cells.

**Figure 4.**
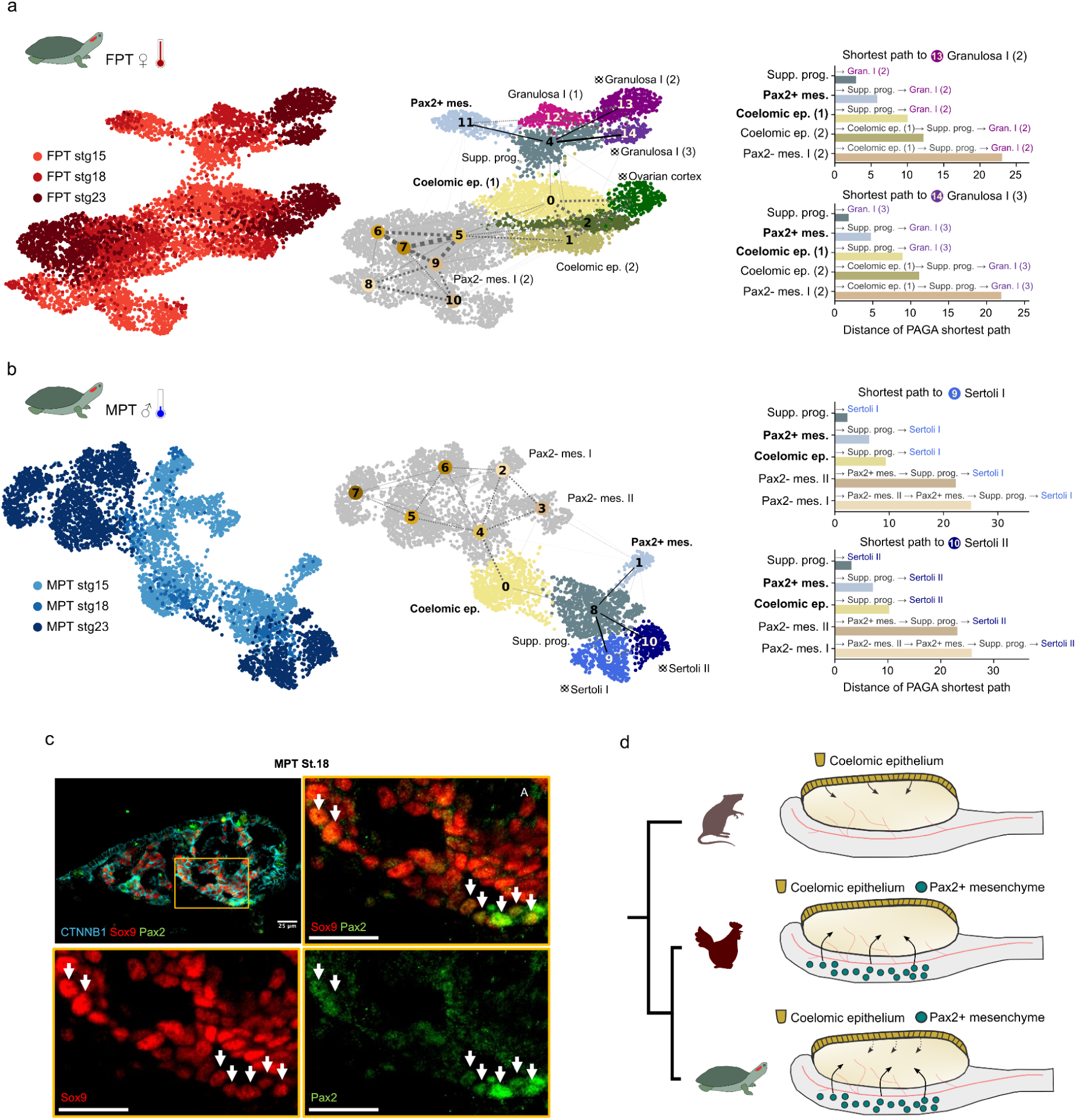
Pax2+ mesenchymal origin of supporting lineage cells in T. scripta. a) PAGA trajectory analysis of FPT somatic gonad cells. On the left, UMAP embedding from a subcluster of FPT somatic cells displaying information of the stage at which they were sampled. In the middle, the same UMAP embedding but this time colored according to the Leiden subclusters. Subclusters composed of a majority of Stg. 23 cells are considered terminal and marked with a squared flag. The widths of the connections between clusters represent the PAGA scores. Higher PAGA scores suggest a more likely transition between two cell types. The shortest path from Pax2+ mesenchymal cells to granulosa cells is represented with black lines, the rest of connections are shown as discontinuous grey lines. On the right, the barplot represents the shortest distances between early clusters and terminal clusters of granulosa cells. b) Same as i a) but for MPT somatic cells. On the right, shortest distances between early clusters and terminal clusters of Sertoli cells. c) Representative immunofluorescence images of MPT gonadal cross sections at St.18. β-catenin (CTNNB1) is labeled in cyan, Sox9 in red, and Pax2 in green. The yellow box depicts the region magnified on the break-out panels. The white arrows point to cells of the gonad which co-express Sox9 and Pax2 (scalebar, 25 μm). d) Diagram showing an evolutionary scenario in which supporting cells of *T. scripta*, similarly to chicken, are mostly derived from Pax2+ mesenchymal cells.

Next, we characterized in detail the transcriptomic changes that drive the differentiation of supporting cells. We calculated pseudotimes and analyzed genes that changed expression along the differentiation trajectories connecting *Pax2*+ mesenchymal cells to 3 different subpopulations of granulosa (Extended Data Fig. 21) and 2 subpopulations of Sertoli cells (Extended Data Fig. 22). We first focused on the top 50 most expressed TFs that are dynamically expressed along pseudotime. The initial point of all trajectories leading to either Sertoli or granulosa cells was similar, involving the participation of common TFs such as *Pax2*, *Tbx1*, *Id1*, *Meis1* or *Nr2f2* whose expression faded later on (Extended Data Fig. 21c, Extended Data Fig. 22c). Importantly, the expression of TFs like *Tbx1* is maintained in the transition from *Pax2*+ mesenchymal cells to supporting progenitors, overlapping with the TFs characteristic of the somatic gonad (i.e. *Gata4* and *Nr5a1*) and supporting progenitors (*Sox9*). These overlaps in expression are consistent with the coexpression of Pax2 and Sox9 shown by immunofluorescence (Fig. 4c) and further strengthen the idea that early *Pax2*+ cells become supporting cells at later stages. After this first wave, TFs that are exclusively of bipotential supporting cells such as *Stat1*, *Irf1* or *Ctbp1* are subsequently expressed. Those TFs coexist with others that are transiently expressed in one of the sexes during the bipotential period, but continue to be expressed in the opposite sex. Examples of the latter are *Dmrt1*, *Sox8*, *Sox9* and *Esr1* which are all expressed at FPT in a narrow temporal window in supporting progenitor cells, but remain expressed throughout Sertoli differentiation at MPT. Conversely, *Ar* is expressed only briefly at MPT but it is maintained at FPT until granulosa cells are fully differentiated. Eventually, sex specific TFs like *Twist1* and *Foxl2* in granulosa cells or *Nkx3-1* at MPT take over.

Based on our observation that the supporting lineage synthesizes sex hormones, we investigated the temporal dynamics of steroidogenic enzyme expression throughout their differentiation. *Star, Cyp17a1* and *Hsd3b* are predominantly expressed in intermediate states of differentiation at FPT, corresponding to bipotential supporting cells (Extended Data Fig. 21c). Meanwhile, *Cyp11a1* is expressed during the whole process of differentiation to granulosa, although more prominently at early states. Finally, *Cyp19a1* is exclusively expressed later on in cells committed to the granulosa fate. Similar to FPT, *Cyp17a1* is detected only in the intermediate part of the path of Sertoli differentiation at MPT (Extended Data Fig. 22c). However, *Cyp19a1* transcripts are never detected at MPT and *Hsd3b* and *Star* expression are maintained in differentiated Sertoli cells. Furthermore, *Cyp11a1* expression is absent from differentiated Sertoli cells. Finally, we explored the expression of a curated set of genes implicated in the differentiation of granulosa and Sertoli cells in mice ^33^.

As expected, most genes implicated in granulosa differentiation in mice were expressed late in the *Pax2*+ mesenchyme to granulosa differentiation trajectory in *T. scripta* (i.e. *Fst*, *Lef1*, *Axin2*) but notable absences were also noticed (i.e. *Rspo1*, *Bmp5*, *Lgr5* and *Irx3* were either not detected or very lowly expressed, Extended Data Fig. 21c). The genes implicated in Sertoli differentiation in mice were also predominantly expressed later on in the *Pax2*+ mesenchyme to Sertoli in *T. scripta,* with some exceptions such as *Fgf9*, which is only transiently expressed in intermediate states of differentiation in both species (Extended Data Fig. 22c). Altogether, our data show that turtle supporting cells originate primarily from *Pax2*+ mesenchymal cells, as in birds. This supports a mesenchymal origin as the ancestral condition in Archelosauria and contrasts with the mammalian model, where supporting cells emerge predominantly from the coelomic epithelium (Fig. 4d).

## DISCUSSION

Sex determination mechanisms evolve rapidly within vertebrates, ranging from environmental sex determining systems like those based on temperature (TSD) to genetic sex determination (GSD) ^1^. Yet, despite stark differences at the top of sex determining cascades, the overall organization of the testis and ovary organs is similar among vertebrates. Consequently, the cell types, developmental processes and underlying genetic programs involved in sex determination might also be evolutionary conserved ^7^. Here we tested this hypothesis by exploring the cell types and transcriptional programs involved in primary sex determination in a species with TSD, the turtle *T. scripta*, using single-cell transcriptomics. We contextualize our results using equivalent datasets from vertebrate species with GSD: chicken and mouse. By focusing on the expression of known markers and by comparing expression programs across species we confirmed an overall conservation of the main gonadal lineages: germinal, supporting and interstitial. Our results suggest that these three lineages are conserved at least in the last common ancestor of amniotes. However, despite the broad conservation of gonadal lineages across amniotes, we found that the distribution of some gonadal functions differs.

An intriguing finding is that *T. scripta* gonads lack fetal Leydig cells, a cell type responsible for androgen production in mammals. The absence of Leydig cells in early turtle gonads is notable, since sex hormones have been repeatedly shown to influence gonadal sex differentiation in *T.scripta* ^19,21^, and Leydig cells have been described in the adult testes of several turtle species ^21^ ^34,35^. However, fetal and adult Leydig cells have a different developmental origin, as reported in mammals ^15^, consistent with the observations that one of these populations is absent in early prenatal turtle gonads. Here, we show that this critical steroidogenic function in fetal turtle gonads is mediated by the supporting lineage, including progenitors and differentiated Sertoli and granulosa cells. Of note, Sertoli cells in the adult testis of the turtle *C. serpentina* also produce androgens required for spermiogenesis ^36^ thus confirming that the adult supporting lineage in turtles is competent for the synthesis of steroid hormones. Outside mammals, fetal Leydig cells have only been described in chicken^10^. However, these cells derive from Sertoli cells, in contrast to mammals, where they differentiate from interstitial progenitors. In fact, steroidogenic activity was detected within the sex cords of chicken gonads, and not only in the interstitium. Therefore, the homology between interstitially-derived Leydig cells in mammals and Sertoli-derived steroidogenic cells in birds remains uncertain. Based on these data, we hypothesize that outside mammals, fetal steroidogenesis is driven by supporting-lineage cells with varying levels of specialization, whereas *bona fide* interstitial fetal Leydig cells may represent a mammalian innovation. Thus, in non-mammalian vertebrates, where sex hormones can have a dominant effect on gonadal sex differentiation, these hormones may be synthesized by the same lineage that initiates sex determination: the supporting lineage.

Recent single-cell studies of sex determination in *T. scripta* identified a population of thermosensitive steroidogenic cells (TSSCs), proposed to initiate testis development through aldosterone secretion ^37,38^. These studies suggest that TSSCs are located within the developing gonad and are independent of the supporting lineage, based on immunofluorescence analyses of putative TSSC markers (Kcnk3, Cacna1h, and Cacna1c). While we also identify this population in our dataset, multiple lines of evidence indicate that these cells are located outside the gonad, and we therefore refer to them as non-gonadal steroidogenic cells. First, immunofluorescence analyses reveal an almost exclusive colocalization of steroidogenic and supporting cell markers (Cyp11a1 and Sox9, respectively) within the gonad, whereas steroidogenic cells lacking supporting markers are mainly located in the adjacent mesonephros (Fig. 3h). Consistently, non-gonadal steroidogenic cells are detected only at early developmental stages, when mesonephric tissue is included in the dissections, and form a distinct population in UMAP space, with no apparent transcriptional continuity with other somatic gonadal lineages (Fig. 1c). Finally, we find that the proposed TSSC markers Kcnk3, Cacna1h, and Cacna1c are not specific to TSSCs, as they are also expressed in supporting cells, both in our dataset and in the cited studies (see Supplementary Figure 3 of Ye et al. ^38^). Thus, these markers cannot reliably resolve the spatial localization of TSSCs. Yet, despite the noted discrepancy, this collective work supports the notion that steroidogenic cell populations, whether intrinsic to the gonad or closely associated with it, contribute to temperature-dependent sex determination in turtles.

The steroidogenic role of the supporting lineage in Archelosauria (turtles and birds) and its contrast with mammals, where these cells do not produce sexual hormones, constitutes an example of functional divergence. These distinct functions may be related to the different developmental origins of these putative homologous cell lineages. As we show, granulosa and Sertoli cells in turtles originate primarily from a Pax2+ mesenchymal population, similar to birds ^10,13^, while most supporting cells arise from the coelomic epithelium in mammals ^12^. We hypothesize that both progenitor pools had the potency to give rise to supporting lineage cells in the last common ancestor of amniotes. Later, either the coelomic epithelium or the Pax2+ mesenchymal cells may have gained importance in each lineage independently. This hypothesis is supported by the recent discovery of supporting-like cells (SLC) in mammalian gonads ^14,32^. SLCs are similar transcriptomically to supporting cells and they mainly contribute to the formation of the rete testis and rete ovarii in males and females respectively, but they also constitute a minor source of Sertoli and granulosa cells. Interestingly, they are characterized by the expression of the TF Pax8, a close paralog of Pax2 (the PaxB group is composed of *Pax2*, *Pax5* and *Pax8*, which originated in the two rounds of whole genome duplications of vertebrates) ^39^. Alternatively, the capacity of the coelomic epithelium to give rise to supporting cells could have appeared in mammals. A denser taxon sampling within amniotes and also beyond, including amphibians where sex hormones are critical ^40^, could help clarify the evolutionary history of supporting lineage cells.

Besides differences in cell-type repertoire and origins, we also found divergence in the genetic programs of conserved cell types, involving important lineage-specific regulators such as *Tbx1* or *Esr1*. The absence of *Runx1* expression in turtle granulosa cells, and of several of its direct downstream targets, highlight how gene regulatory networks may reconfigure upon the loss of critical TFs ^31^. Particularly striking was the discovery of Twist1 as the earliest TF acquiring sexual dimorphism during turtle ovarian differentiation, prior to the onset of Foxl2 expression. While Twist1 has never been associated with sex determination previously, it has been shown to act as a repressor of the male sex determining gene Sox9 in the context of chondrogenesis ^24^. Importantly, Twist1 has been shown to be activated by pStat3 in cancer cell lines ^41^ and the phosphorylation of Stat3 in response to high temperatures is one of the first known events in ovarian differentiation in *T. scripta*. pStat3 was shown to promote ovarian differentiation due to the indirect repression of *Dmrt1* via the repression of the Kdm6b demethylase ^20^. Here we propose that, additionally, pStat3 could promote ovarian differentiation by activating the expression of the female-specific TF Twist1. Importantly, pStat3 has also been implicated in the direct activation of the classic female promoting TF Foxl2 ^42^, yet our single cell datasets suggest that the expression of *Twist1* predates *Foxl2* expression in the sex determining cascade of turtles. *Twist1* promotes epithelial to mesenchymal transition (EMT) in embryonic ^43^ and cancer ^44^ cell types, which could be important for the reorganization of the primary sex cords in the medulla of the turtle ovary. Overall we found that cell type repertoires, developmental processes and genetic programs involved in vertebrate sex determination are more divergent than previously anticipated. This divergence parallels, to some extent, the turnover of sex determination systems, offering ample regulatory flexibility that may be crucial for species adaptation to changing environments.

## METHODS

### Turtle eggs sampling

Freshly laid red-eared slider turtle (*Trachemys scripta elegans*) eggs were obtained from Concordia Turtle Farms (Jonesville, LA) with permission from the Louisiana Department of Agriculture. One day after oviposition, fertilized eggs were randomly distributed into trays of moist perlite and incubated at either 26 °C (male-producing temperature) or 31 °C (female-producing temperature), with humidity maintained at 70–80%. To minimize clutch effects, eggs from the same clutches were allocated across treatments. Embryos were staged according to the criteria established by Greenbaum ^45^. For all experiments, embryos were removed from the eggshells, immediately decapitated, and transferred to sterile PBS for gonad dissection.

### Single-cell RNA-seq library preparation

Eight to twelve pairs of gonads per temperature and stage were pooled for each single-cell RNA-seq experiment. Cell suspensions were prepared using 0.05% Trypsin-EDTA (59417C, Sigma) for 7 min at either MPT or FPT, depending on the sample’s original incubation condition. During digestion, tissue was gently pipetted every 2–3 min. The reaction was stopped by adding 5% BSA (B9000S, New England Biolabs), and the suspension was passed through a 40 µm Flowmi cell strainer (15342931, Fisher Scientific). Cells were then collected by centrifugation (300 × g for 5 min at 4 °C), resuspended in PBS for counting and viability assessment, and subsequently fixed by adding methanol (98%) dropwise. Fixed cells were kept on ice for 15 min before being transferred to −80 °C for long-term storage. After rehydration, about 10,000–20,000 cells were loaded on a 10× Chromium controller and sc-RNA libraries were prepared using the Chromium Next Gem Single Cell 3’ Reagent Kit v3.1 (PN-1000128) following the manufacturer’s instructions. Finally, libraries were sequenced using Illumina NovaSeq 6000 SP with the following parameters: paired-end, 28 + 8 + 91 cycles. The sequencing depth was 200 million reads per sample.

### Single-cell RNA-seq analysis

Each *T. scripta* single-cell library was aligned to the *T. scripta* assembly and annotation (CAS_Tse_1.0, GCA_013100865.1) using Cell Ranger. The three prime ends of the genes of the annotation were corrected and extended to improve mapping. Quality control of the cells was performed considering the number of reads and features detected, the percentage of mitochondrial transcripts, and ambient RNA contamination with DecontX^46^. For chicken and mouse, filtered expression matrices were downloaded from GEO (see Data Availability). Doublets were filtered out using scrublet^47^ in all cases. Cells with low fraction of nuclear mRNAs were also filtered out in *T. scripta* and chicken datasets with DropletQC^48^. The different samples within each species were integrated using SCVI^49^ (https://scvi-tools.org/) and SCVI models were also used for differential expression analysis. The number of latent variables was optimized for both the full gonad by doing a parameter sweep and inspecting the learning curves. Both UMAP dimensionality reductions and leiden clusters were calculated from SCVI latent variables using scanpy^50^. Marker genes for each of the clusters were obtained using SCVI differential expression of each cluster against the rest. GSEA analyses were run on the obtained fold changes of genes that had 1 to 1 orthologs with mouse using clusterProfiler^51^. Combined scores for particular sets of genes such as genes from a particular GO term were calculated with AUCell^52^. WGCNA module detection and cross-species comparisons were performed with the package hdWGCNA^53^ following the recommendation of the vignette to choose parameters. Only 1 to 1 orthologs between the species to compare were used both to calculate and to compare the modules. NetRep^54^ was used to assess the conservation of the WGCNA modules, which provide both density and connectivity metrics. In short, density metrics assess whether the genes of the module are also coexpressed in a second species, while connectivity metrics measure whether the hierarchies within the module are maintained. Therefore, we always focus first on the conservation of density metrics. The list of 1 to 1 orthologs was refined by using TOGA ^55^ between *T. scripta* and chicken. For trajectory inference, new SCVI models were created using only cells from male or female samples and cell types from the somatic gonad and the mesonephros (excluding germ cells, endothelial cells, kidney, etc.) After reclustering these subsets of gonadal cells, PAGA^56^ was applied to calculate likely transitions between clusters. Then, PAGA connectivities were converted to distances and Dijkstra’s algorithm was used to calculate the minimal distance between clusters composed of cells from early stages to cells from later stages. Diffusion components were calculated using destiny ^57^ and the pseudotime of the different lineages with slingshot ^58^. Genes that were dynamically expressed along the pseudotime and the smoothed expression values were obtained with tradeSeq ^59^. Custom R and python code to reproduce the analyses and the figures is available in GitLab (https://gitlab.com/rdacemel/trachemys-sd) and contains custom python and R functions gathered in two packages under development, bacarisas (https://gitlab.com/rdacemel/bacarisas, for python) and sorolla (https://gitlab.com/rdacemel/sorolla, for R) which focus in the production of custom plots.

### Immunostaining of cryosections

Gonad–mesonephros complexes (GMCs) and whole gonads were dissected in PBS and fixed in 4% PFA (AAJ19943K2, Fisher Scientific) for 30 min rocking at room temperature, then moved through a MeOH gradient to 100% MeOH for long-term storage at −20 °C. Samples for cryosection were gradually rehydrated into PBS, moved through a sucrose gradient (10%, 15%, 20%, and 30%), then placed in blocks with OCT medium and moved to −80 °C for a minimum of 12 h before cryosectioning. Blocks were serially sectioned at 12 μm and slides were stored at −20 °C until ready to use. For immunostaining, sections were rehydrated in PBS, permeabilized in PBST (0.1% Triton X-100), and blocked for 1 h at room temperature (PBS 0.1% TritonX-100, 3% BSA, 10% horse serum). Sections were incubated in primary antibodies diluted in blocking solution at 4 °C overnight (Extended Data Table 1). Following three 10-min washes in PBS 0.1% TritonX-100, sections were incubated with secondary antibodies (Cy3 Donkey Anti-rabbit Jackson Immuno Cat# 711-165-152; AF488 Donkey Anti-mouse Jackson Immuno Cat# 715-545-150; AF647 Donkey Anti-goat Jackson Immuno Cat# 705-605-147) at 1:500 dilution and DAPI (62248,Thermo Fisher) diluted in blocking solution for 2 h at room temperature. Following three 10-minute washes in PBST, slides were mounted for imaging with polyvinyl alcohol mounting solution (Catalog #) and stored at 4 °C. Sections were imaged using Leica SP8 confocal microscope and images were analyzed using FIJI ^60^.

## DATA AVAILABILITY STATEMENT

Single cell RNA-seq (scRNA-seq) data in *T. script*a have been deposited in GEO repository (GSE271230). Equivalent single cell datasets in chicken and mouse are available under the accessions GSE143337 and GSE184708 respectively.

## CODE AVAILABILITY

The custom R and python code to reproduce the analyses and the figures is available in GitLab (https://gitlab.com/rdacemel/single-cell-TSD) and contains custom python and R functions gathered in two packages: *bacarisas* (https://gitlab.com/rdacemel/bacarisas, for python) and *sorolla* (https://gitlab.com/rdacemel/sorolla, for R) which focus in the production of custom plots.

## AUTHOR CONTRIBUTIONS

R.D.A, B.T., B.C., and D.G.L. conceived the study and designed experiments. B.T. collected and preprocessed Trachemys gonads with the help of C.W.. R.D.A., W.Y.C. and A.H. performed the single-cell RNA-seq experiments. B.T. and S.A. performed immunostainings. R.D.A. analyzed data with inputs from B.T., B.C. and D.G.L.. R.D.A, B.T, B.C and D.G.L wrote the paper with input from all authors.

## ACKNOWLEDGEMENTS

We thank the sequencing core of the Max Delbrück Centre for Molecular for technical assistance and the members of the Lupiáñez, Capel, and Tezak Labs for thoughtful discussions and feedback. We are also grateful to the Duke Light Microscopy Core Facility (NIH Shared Instrumentation Grant 1S10RR027867) and the Wesleyan Advanced Microscopy Core Facility. Research in the Lupiañez lab was funded by the European Research Council (grant no. 101045439, 3D-REVOLUTION) and by the Spanish “Agencia Estatal de Investigación” (grant number PID2022-143253NB-I00/ AEI/10.13039/501100011033/ FEDER, UE). Funded by the European Union. Views and opinions expressed are however those of the author(s) only and do not necessarily reflect those of the European Union or the European Research Council Executive Agency. Neither the European Union nor the granting authority can be held responsible for them. R.D.A. was supported by an EMBO Postdoctoral Fellowship (Grant no. EMBO ALTF 537-2020) and by the “Agencia Estatal de Investigación” (Ramón y Cajal RYC2023-045620-I). We also acknowledge support by the NIH Grant R01HD103064 and NSF Grant 1854642 to B.C.

## EXTENDED DATA TABLES

**Extended Data Table 1.** Reference of antibodies used in immunofluorescence assays.

| Antibody | Raised in | Working dilution | Vendor | Cat. No. |
| --- | --- | --- | --- | --- |
| Gata4 | Goat | 1:500 | Santa Cruz | SC-1237 |
| Gata4 | Mouse | 1:500 | Santa Cruz | Sc-25310 |
| HuC/ HuD | Mouse | 1:50 | Thermo Fisher | A-21271 |
| Sox9 | Goat | 1:500 | R&D Systems | AF3075 |
| Lhx9 | Goat | 1:200 | Santa Cruz | SC-19348 |
| Twist1 | Rabbit | 1:100 | Cell Signaling | 90445 |
| CTNNB1 | Mouse | 1:200 | Sigma Aldrich | C7207 |
| Laminin | Rabbit | 1:500 | Gift from Erikson lab (Duke University) | N/A |
| Pax2/8 | Rabbit | 1:250 | Thermo Fisher | 10336-1-ap |
| Cyp11a1 | Rabbit | 1:250 | Novus | Nbp1-85368 |

## Extended Data Figures for

**Extended Data Figure 1:**
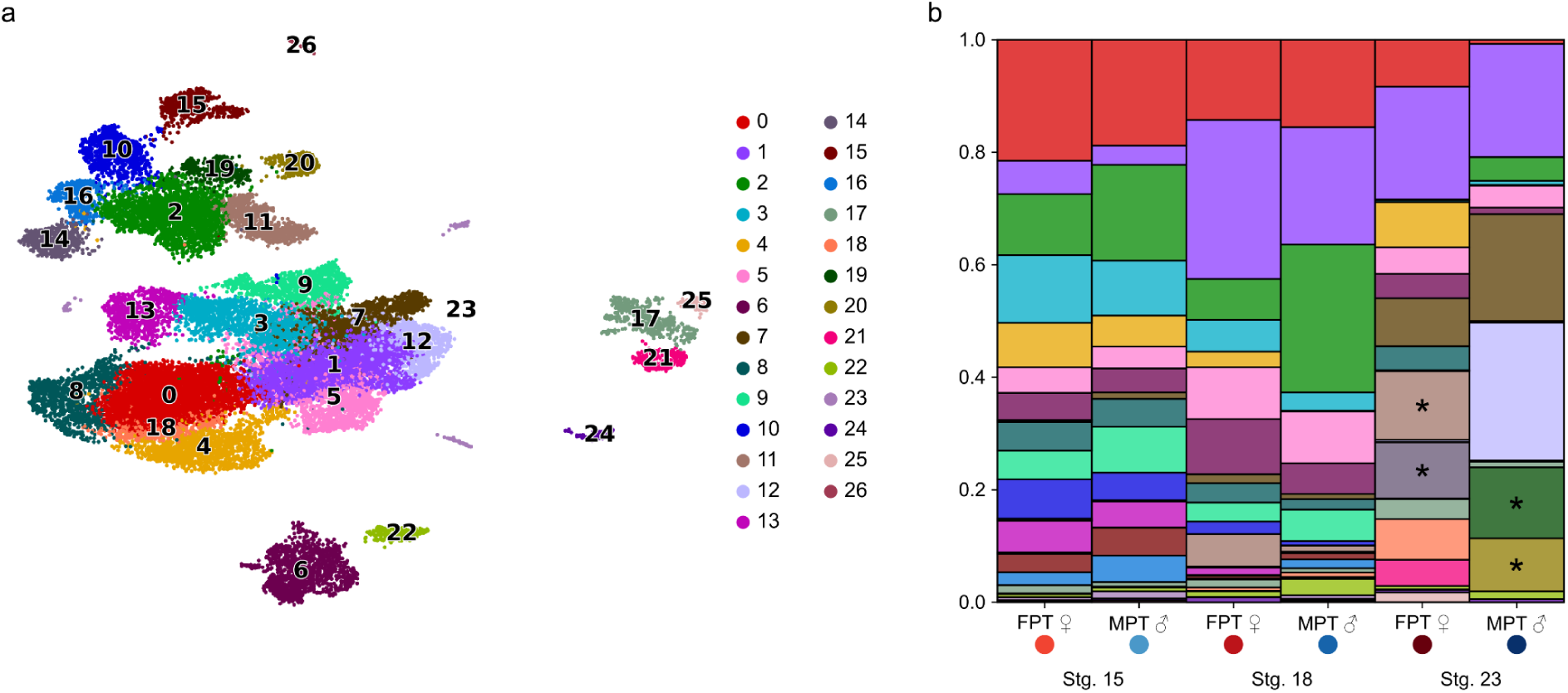
The cell types of *T. scripta* gonads. a) UMAP projection after the integration of the six different samples colored by clusters calculated via Leiden. b) Stacked barplot with the cell type composition of each of the 6 different samples. Colors as in a). Note the similar compositions in Stg.15 samples and the sex-specific clusters in Stg. 23 samples (asterisks).

**Extended Data Figure 2:**
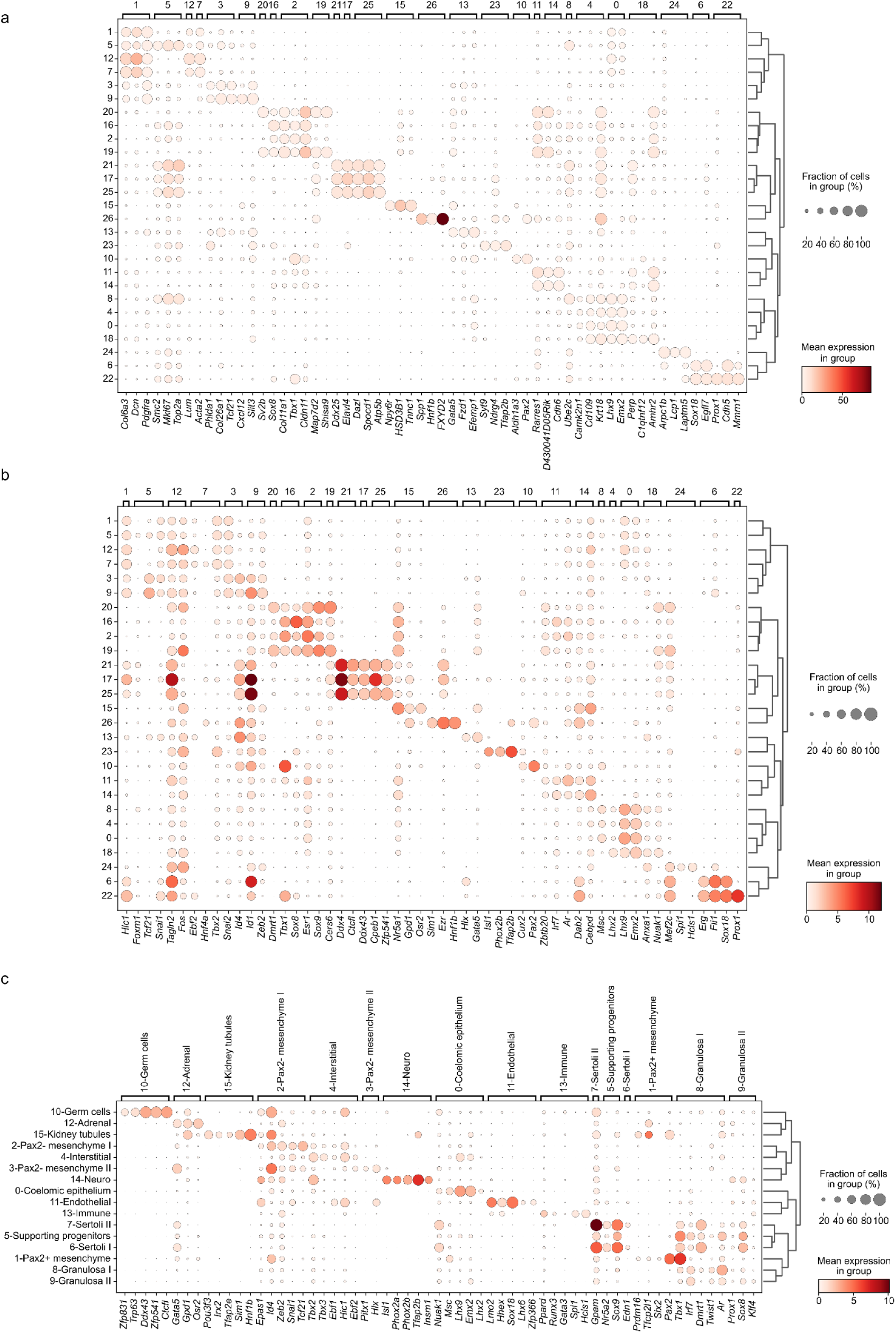
Marker genes and marker TFs of the cell types of *T. scripta* gonads. a) Dotplot displaying the expression of the top 3 non-redundant markers per leiden cluster. The size of the dot represents the percentage of the cells expressing the gene in each cell type and the color the mean expression. b) As in a) but for the top 3 TFs. c) As in b), top 3 TFs per cell type.

**Extended Data Figure 3:**
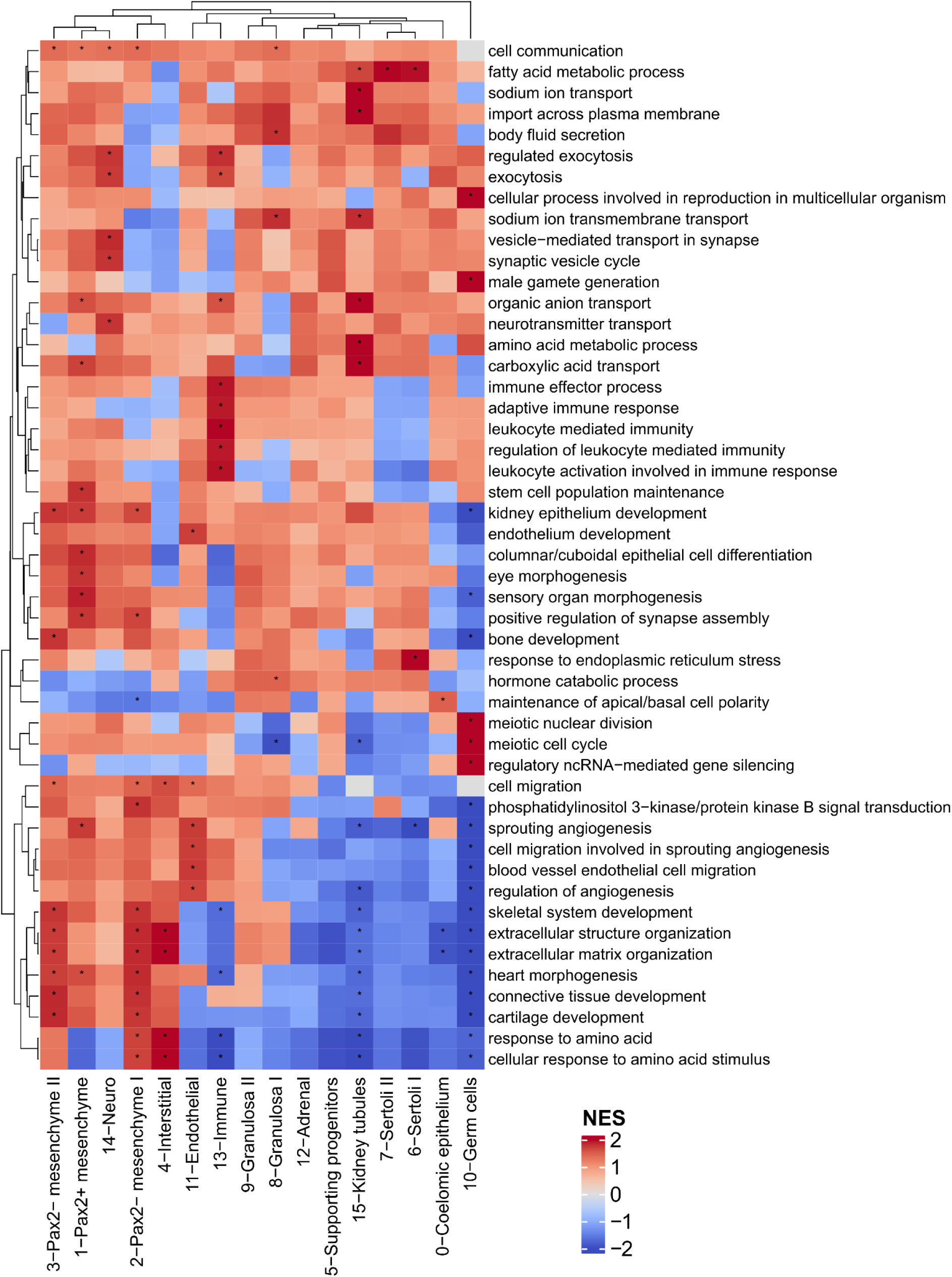
Functional enrichment of the cell types of *T. scripta* gonads. Heatmap displaying Normalized Enrichment Scores (NES) of GO BP terms in the different cell types of *T. scripta* after GSEA analysis. High NES values imply that the genes from a particular GO term are among the highest expressed genes in a particular cell type and viceversa. Asterisks imply significant enrichment or depletion after Benjamini-Hochberg correction.

**Extended Data Figure 4:**
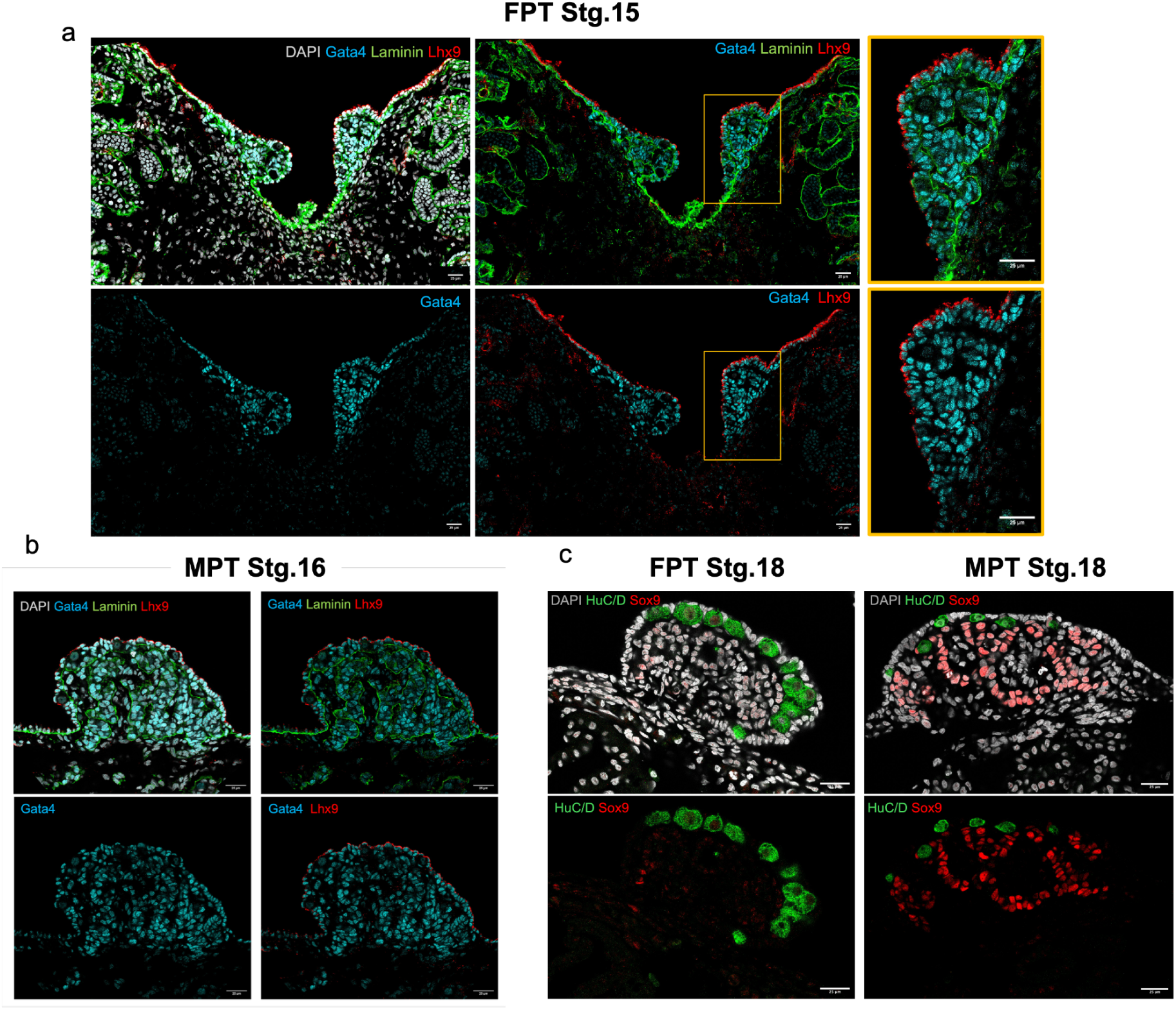
Immunofluorescence localization and validation of single-cell markers. a-b). Representative immunofluorescence images of Stg.15 gonadal-mesonephros complex (a) and Stg. 16 gonads (b). DAPI in grey labels all cell nuclei, Gata4 in Cyan labels gonadal cells, Laminin in green elucidates morphological characteristics of early gonad (and mesonephros), Lhx9 in red labels pluripotent cells from in the coelomic epithelium. c) Representative images of cross sections of FPT and MPT gonads during sex determination stage. DAPI in grey labels all cell nuclei, Germ Cells are labeled in green with HuC/D and supporting cells in red with Sox9 (scalebar, 25 μm).

**Extended Data Figure 5:**
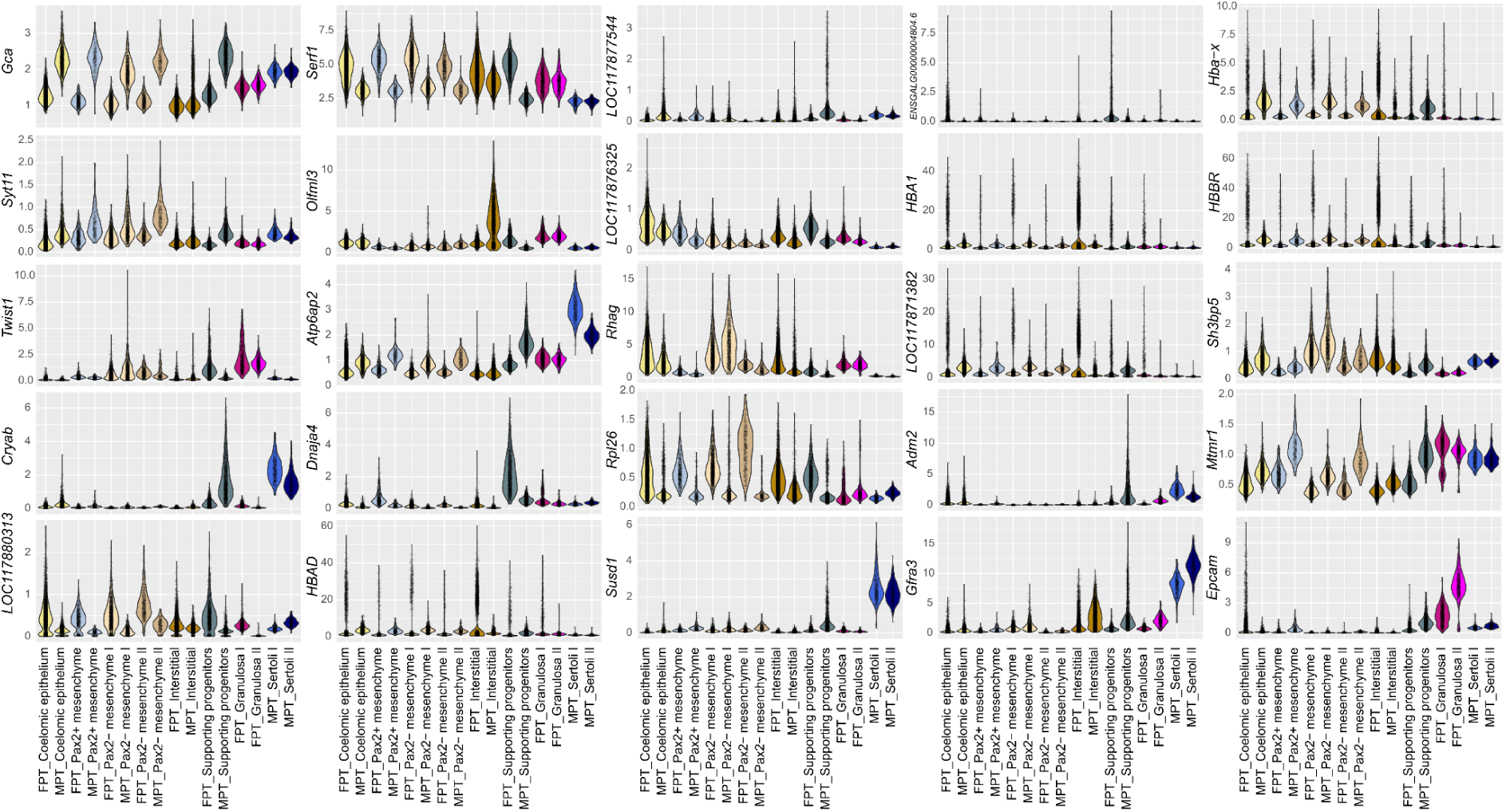
Expression of DEGs in the Supporting progenitors FPT vs MPT comparison. SCVI normalized expression per cluster and per sex of the 25 differentially expressed genes in the FPT vs MPT comparison among supporting progenitor cells.

**Extended Data Figure 6:**
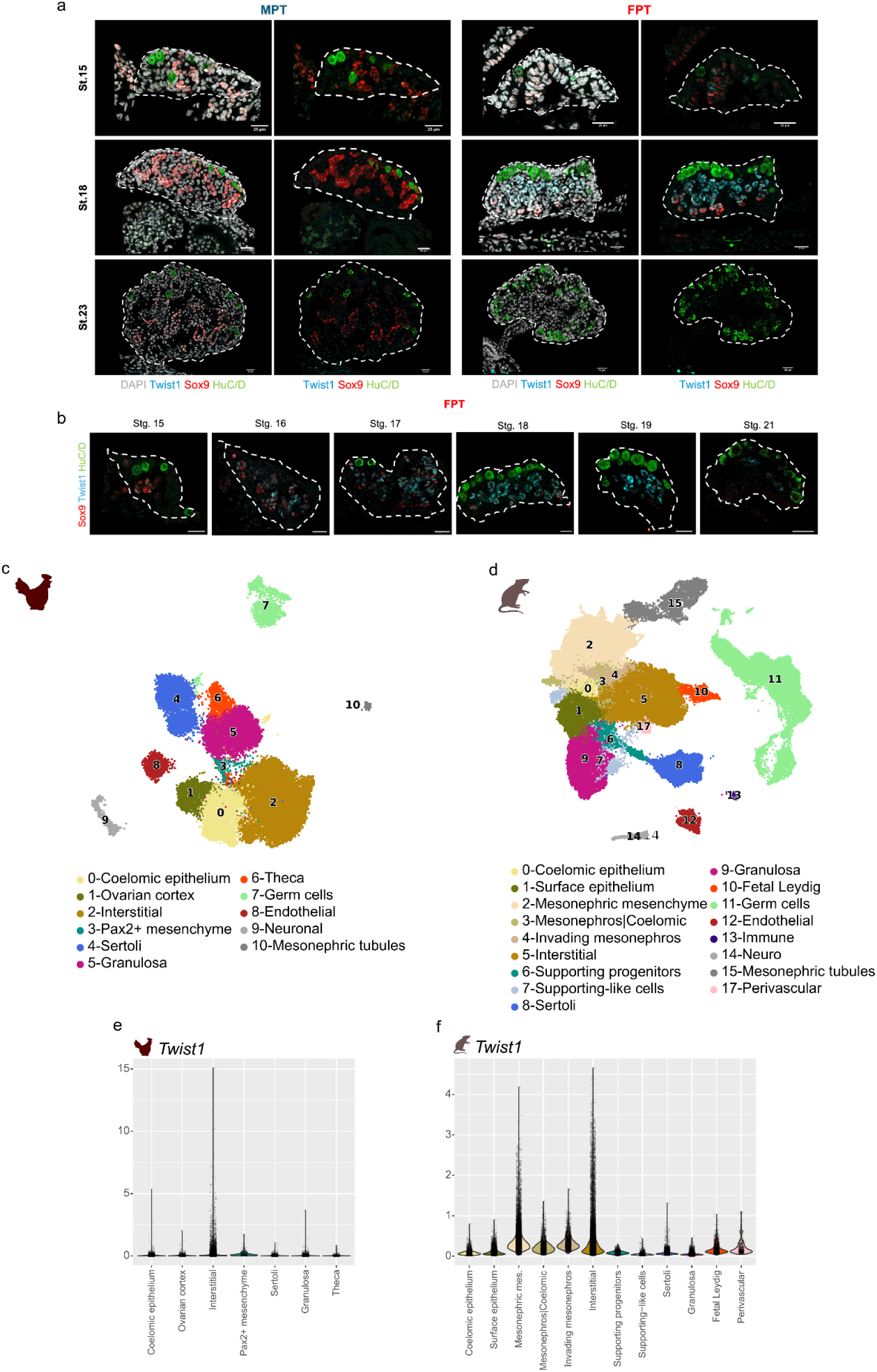
Twist1 expression in vertebrate gonads. a) Comparison of Twist1 and Sox9 expression at MPT and FPT at stgs.15,18 and 23 via Immunofluorescence. Twist1 is labeled in cyan, Sox9 in red, and HuC/D (labelling germ cells) in green. The gonad is outlined with white dashed line. Note the absence of Twist1+ cells at MPT (scalebar, 25 μm) b) Immunofluorescence analysis of Twsit1 and Sox9 expression in FPT gonadal sections throughout sex determination Immuno. Twist1 is labeled in cyan, Sox9 in red, and HuC/D (labelling germ cells) in green. The gonad is outlined with white dashed line (scalebar, 25 μm). c) UMAP projection of SCVI integrated samples of chicken embryonic gonads from Estermann et al. 2020. d) UMAP projection of SCVI integrated samples of mouse embryonic gonads from Mayère et al. 2022. e-f) SCVI normalized expression of *Twist1* in chicken and mouse gonadal cell types.

**Extended Data Figure 7:**
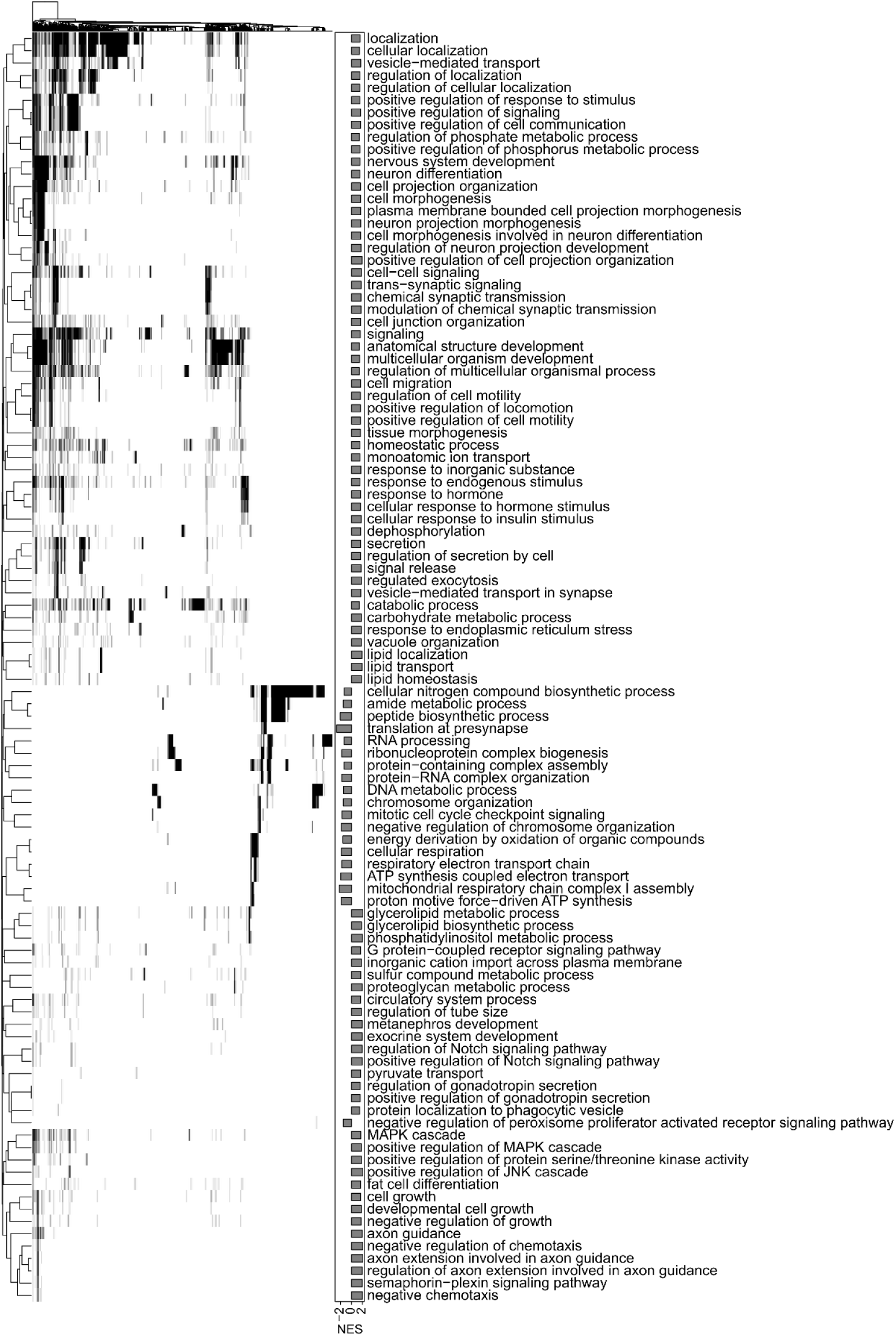
Functional enrichment in the Supporting progenitors FPT vs MPT comparison. Heatmap in which each row represents a GO BP term that is significantly enriched in either FPT (low NES values) or MPT (high NES values). Each column represents a gene. Terms are clustered according to their gene content. If a gene belongs to a particular term the cell will be colored black so that terms that contain similar genes can be appreciated.

**Extended Data Figure 8:**
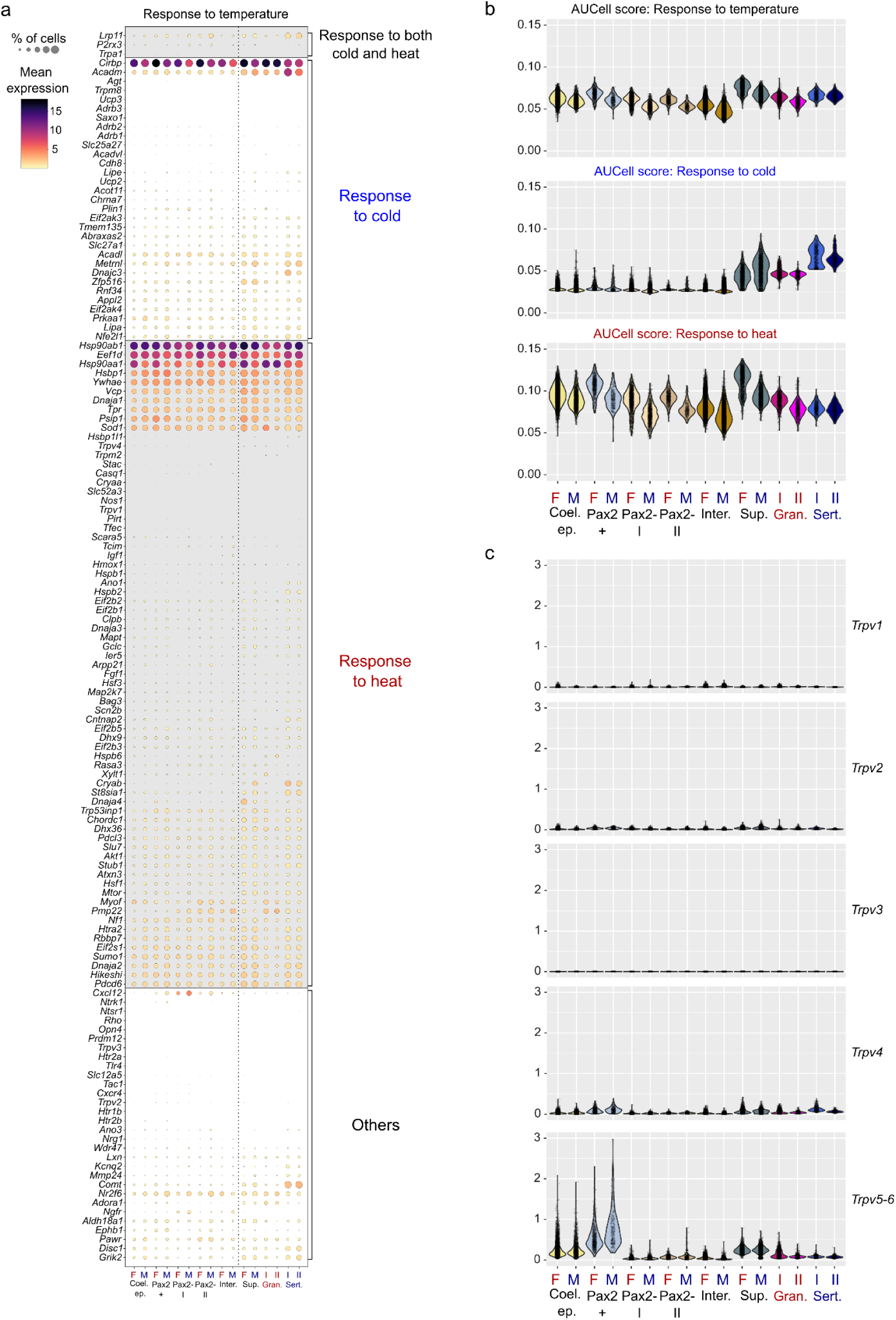
Expression of temperature response genes in *T. scripta* gonads. Dotplot displaying the expression of all the genes belonging to the GO Term *response to temperature.* They are grouped by the subterms *response to heat* or *response to cold*. The size of the dot represents the percentage of the cells expressing the gene in each cell type and the color the mean expression. b) Combined AUCell scores of the gene set composed by the genes belonging to the GO terms in a) in each cell type at either FPT (F) or MPT (M). c) SCVI normalized expression of the genes encoding for the channels of the Trpv family in each cell type at FPT (F) or MPT (M).

**Extended Data Figure 9:**
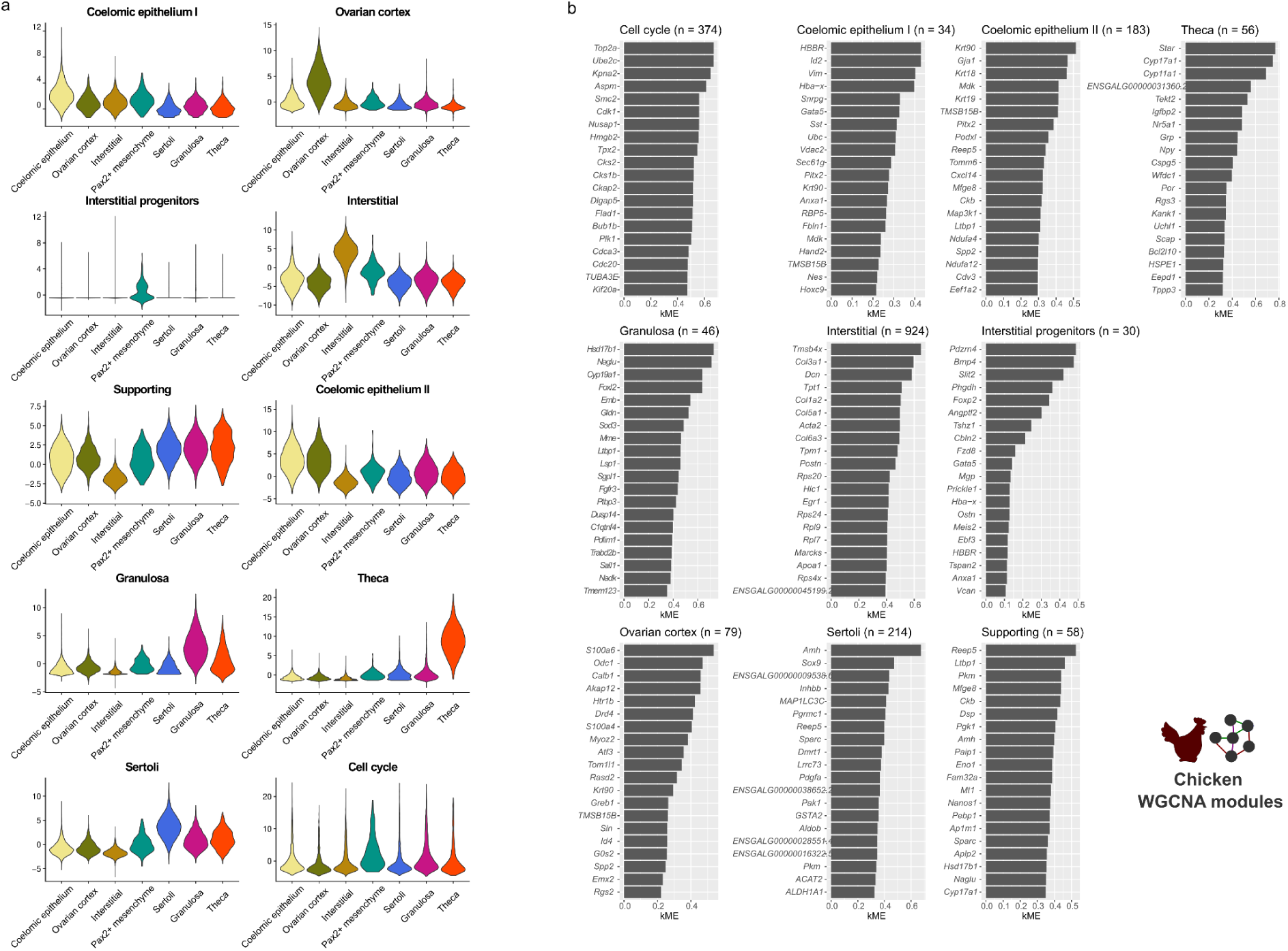
WGCNA modules calculated in the somatic gonad of chicken. a) Violin plot showing the distribution of Module Eigengene (ME) scores of the WGCNA modules in each of the cell types of the somatic gonad of chickens. b) Bar plots showing the membership (kME) of the top 20 genes more associated with each WGCNA module.

**Extended Data Figure 10:**
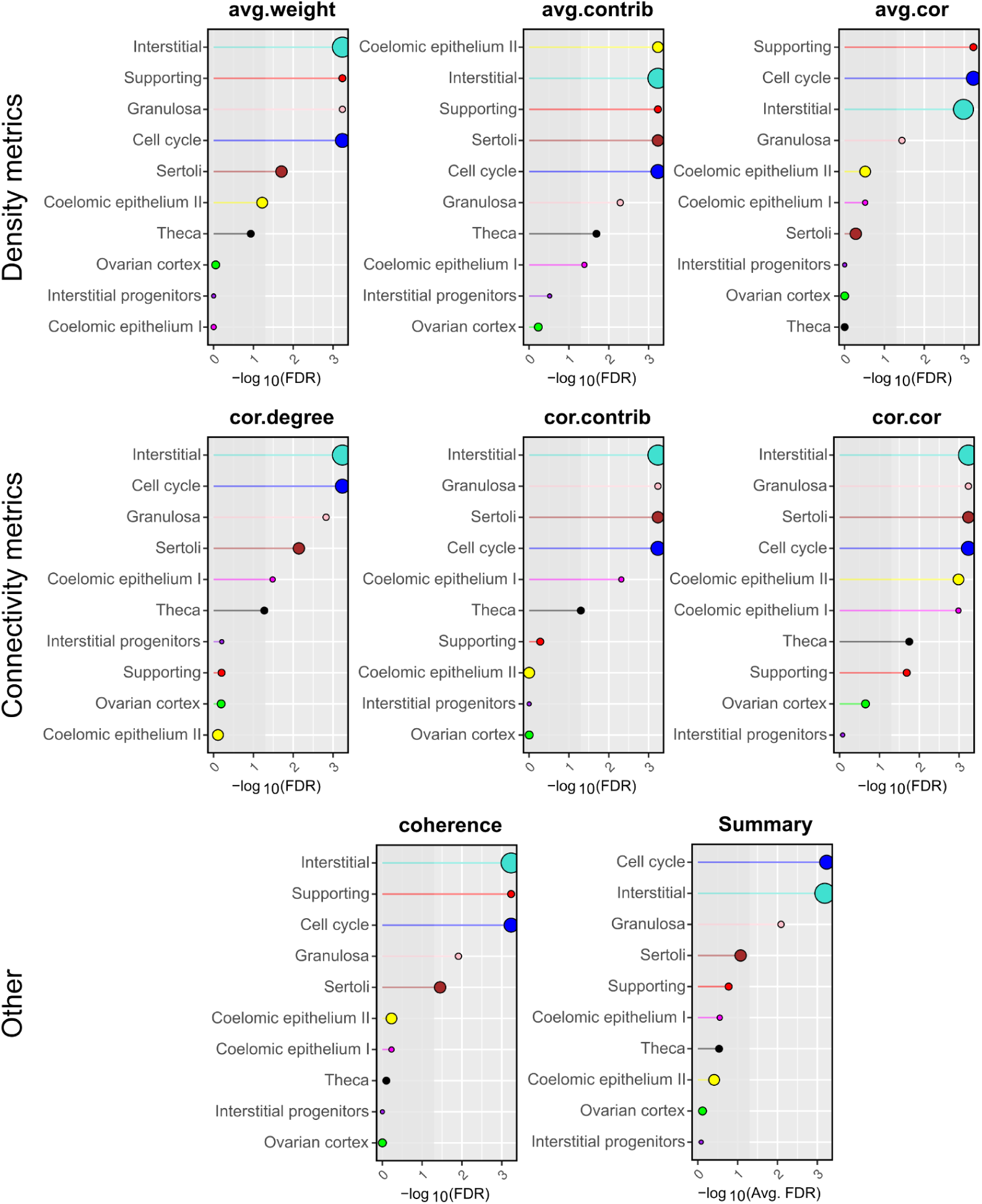
Conservation of chicken WGCNA modules in *T. scripta*. Lollipop plots displaying network conservation FDR values between chicken and *T. scripta* of the different WGCNA modules calculated with NetRep. On the x-axis is represented the negative logarithm of the FDR of a permutation test assessing whether the module has a higher conservation metric than modules generated by chance. Density conservation metrics (top) assess whether the genes that are coexpressed in the WGCNA module in chicken are also coexpressed in *T. scripta.* **avg.weight**: *average magnitude of edge weights*. **avg.contrib**: *average magnitude of the node contribution*. **avg.cor**: *average magnitude of the correlation coefficients*. Connectivity preservation metrics (center) assess whether the hierarchy of the module is conserved, e.g the genes more central to the module in chicken are also central in *T. scripta*. **cor.degree**: *concordance of the weighted degree*. **cor.contrib**: *concordance of the node contribution.* **cor.cor**: *concordance of the correlation structure*. Other metrics like **coherence** measures the variance of the expression in *T. scripta* explained by the WGCNA module in chicken. **Summary** represents the average of all the other metrics. The size of the lollipop circle is proportional to the number of genes in the module.

**Extended Data Figure 11:**
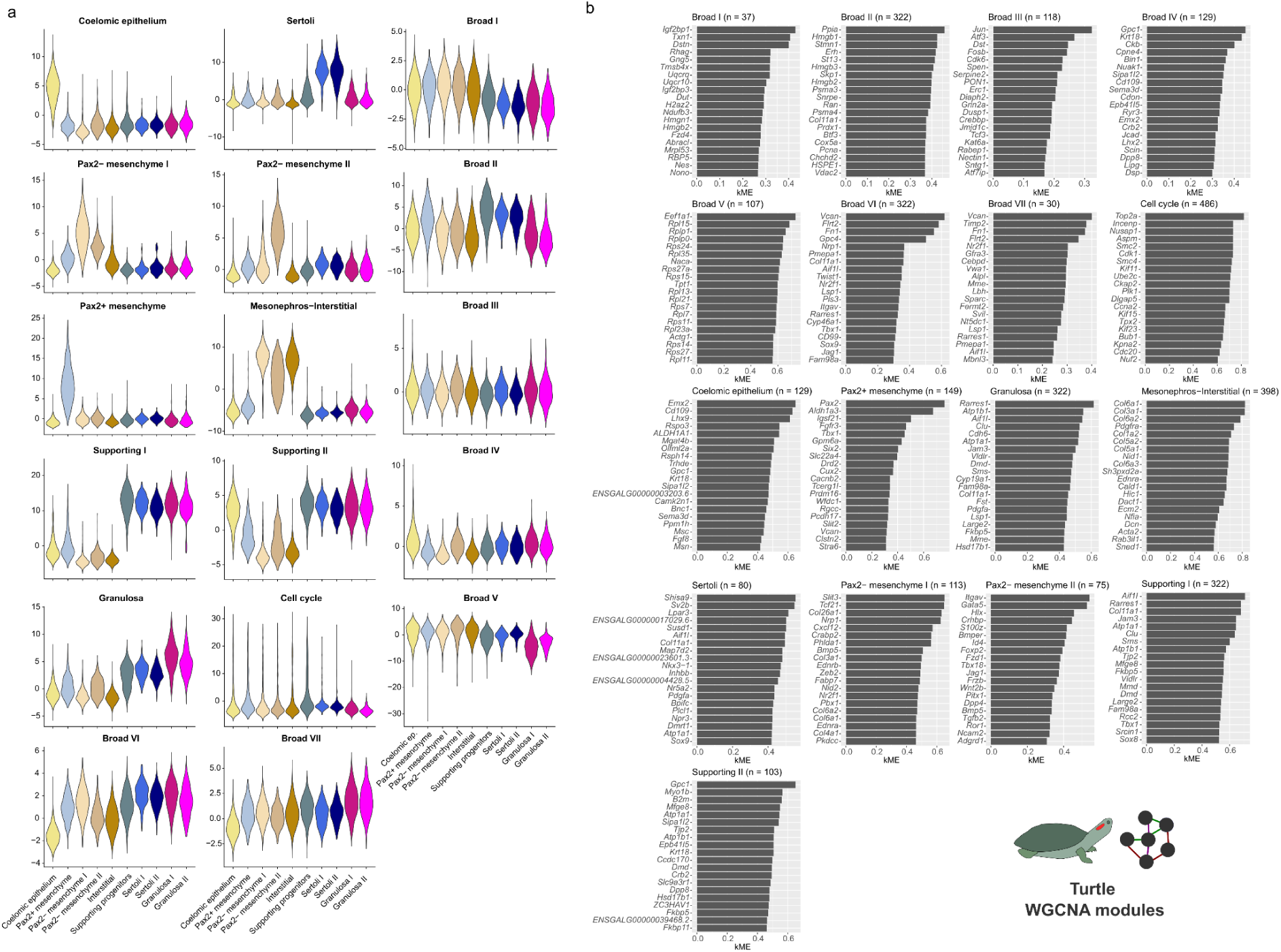
WGCNA modules calculated in the somatic gonad of *T. scripta*. a) Violin plot showing the distribution of Module Eigengene (ME) scores of the WGCNA modules in each of the cell types of the somatic gonad of *T. scripta*. b) Bar plots showing the membership (kME) of the top 20 genes more associated with each WGCNA module.

**Extended Data Figure 12:**
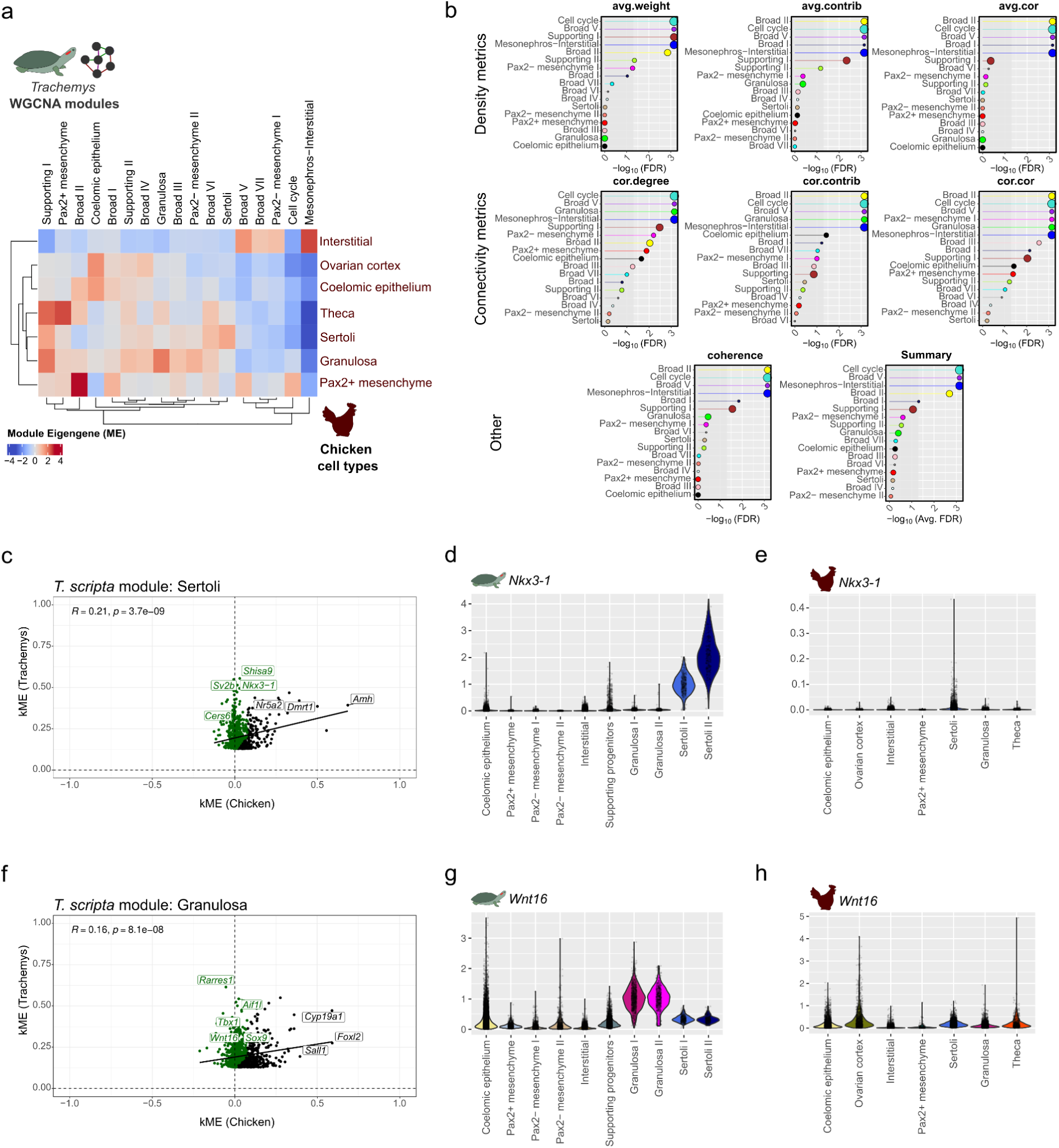
Conservation of *T. scripta* WGCNA modules in chicken. a) Heatmap showing Module Eigengene (ME) scores of *T. scripta* WGCNA modules in each of the cell types of chicken. High ME scores imply that the genes of the *T. scripta* WGCNA module are active in the corresponding cell type of chicken. b) Lollipop plots displaying network conservation FDR values between *T. scripta* and chicken of the different WGCNA modules calculated with NetRep. On the x-axis is represented the negative logarithm of the FDR of a permutation test assessing whether the module has a higher conservation metric than modules generated by chance. See Extended Data Fig. 10. c) Scatterplot showing the membership (kME) of the genes belonging to the WGNCA module *Sertoli* calculated in *T. scripta* both in *T. scripta* (y-axis) and chicken (x-axis). Genes that are associated in *T. scripta* but not in chicken are labeled green (i.e. *Nkx3.1*). d-e) SCVI normalized expression of *Nkx3-1* in *T. scripta* and chicken gonadal cell types. f) As in c) but for the *T. scripta* module *Granulosa*. g-h) SCVI normalized expression of *Wnt16* in *T. scripta* and chicken gonadal cell types.

**Extended Data Figure 13:**
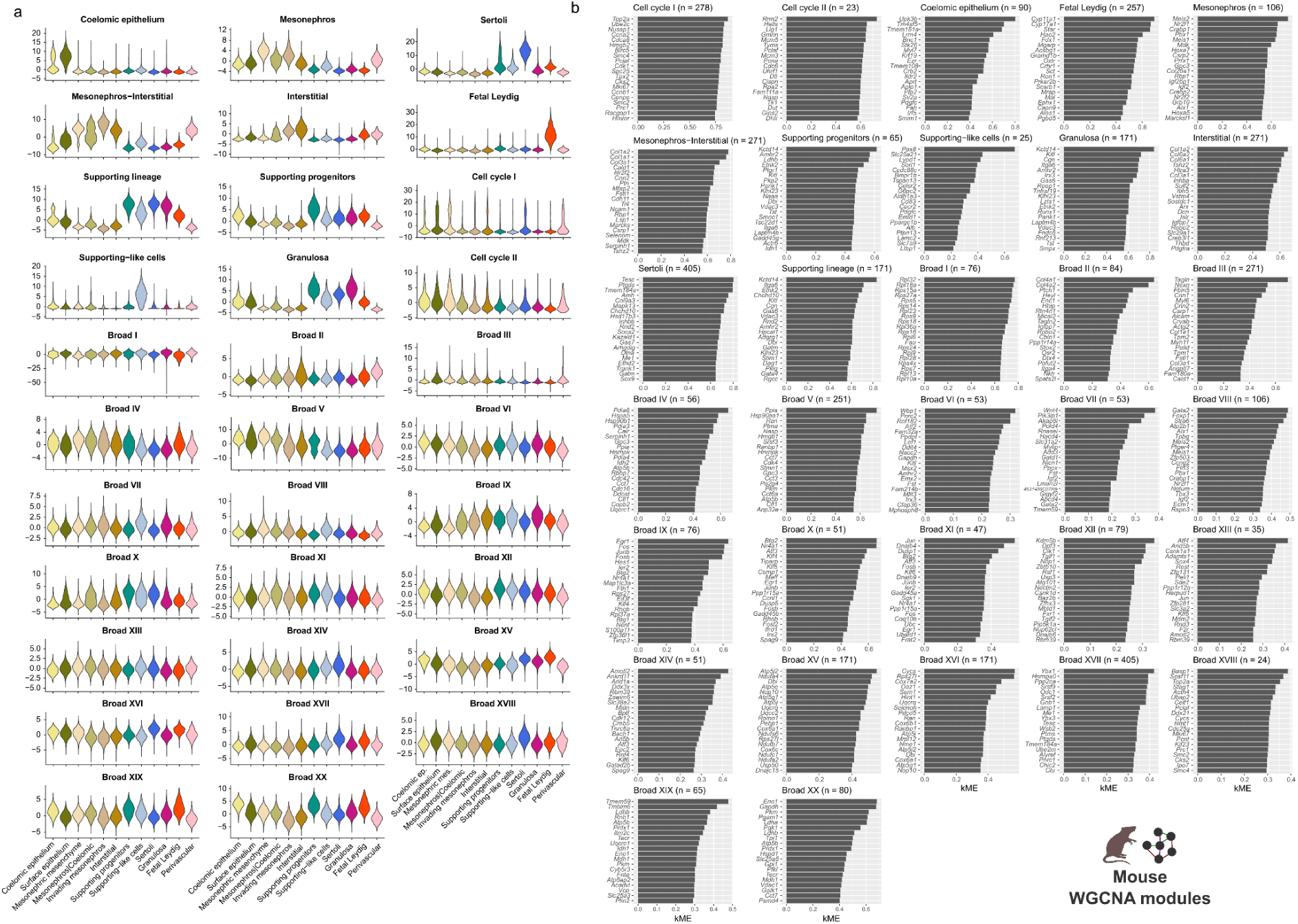
WGCNA modules calculated in the somatic gonad of mouse. a) Violin plot showing the distribution of Module Eigengene (ME) scores of the WGCNA modules in each of the cell types of the somatic gonad of mouse. b) Bar plots showing the membership (kME) of the top 20 genes more associated with each WGCNA module.

**Extended Data Figure 14:**
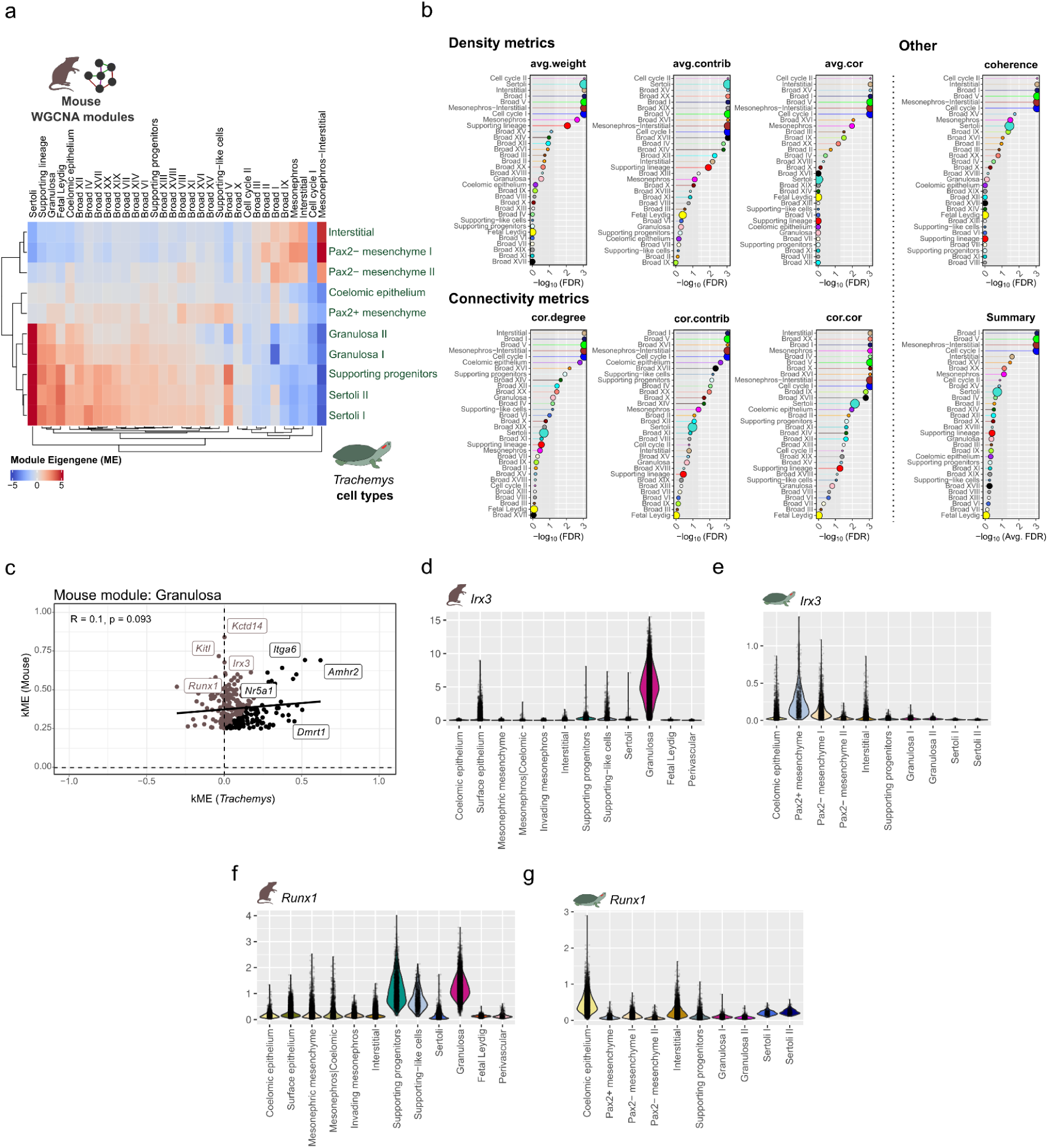
Conservation of mouse WGCNA modules in *T. scripta*. a) Heatmap showing Module Eigengene (ME) scores of mouse WGCNA modules in each of the *T. scripta* cell types. High ME scores imply that the genes of the chicken WGCNA module are active in the corresponding cell type of *T. scripta*. b) Lollipop plots displaying network conservation FDR values between chicken and *T. scripta* of the different WGCNA modules calculated with NetRep. On the x-axis is represented the negative logarithm of the FDR of a permutation test assessing whether the module has a higher conservation metric than modules generated by chance. See Extended Data Fig. 10. c) Scatterplot showing the membership (kME) of the genes belonging to the WGNCA module *Granulosa* calculated in mouse both in mouse (y-axis) and *T. scripta* (x-axis). Genes that are associated in mouse but not in *T. scripta* are labeled brown (i.e. *Irx3* and *Runx1*). d-e) SCVI normalized expression of *Irx3* in mouse and *T. scripta* gonadal cell types. f-g) SCVI normalized expression of *Runx1* in mouse and *T. scripta* gonadal cell types.

**Extended Data Figure 15:**
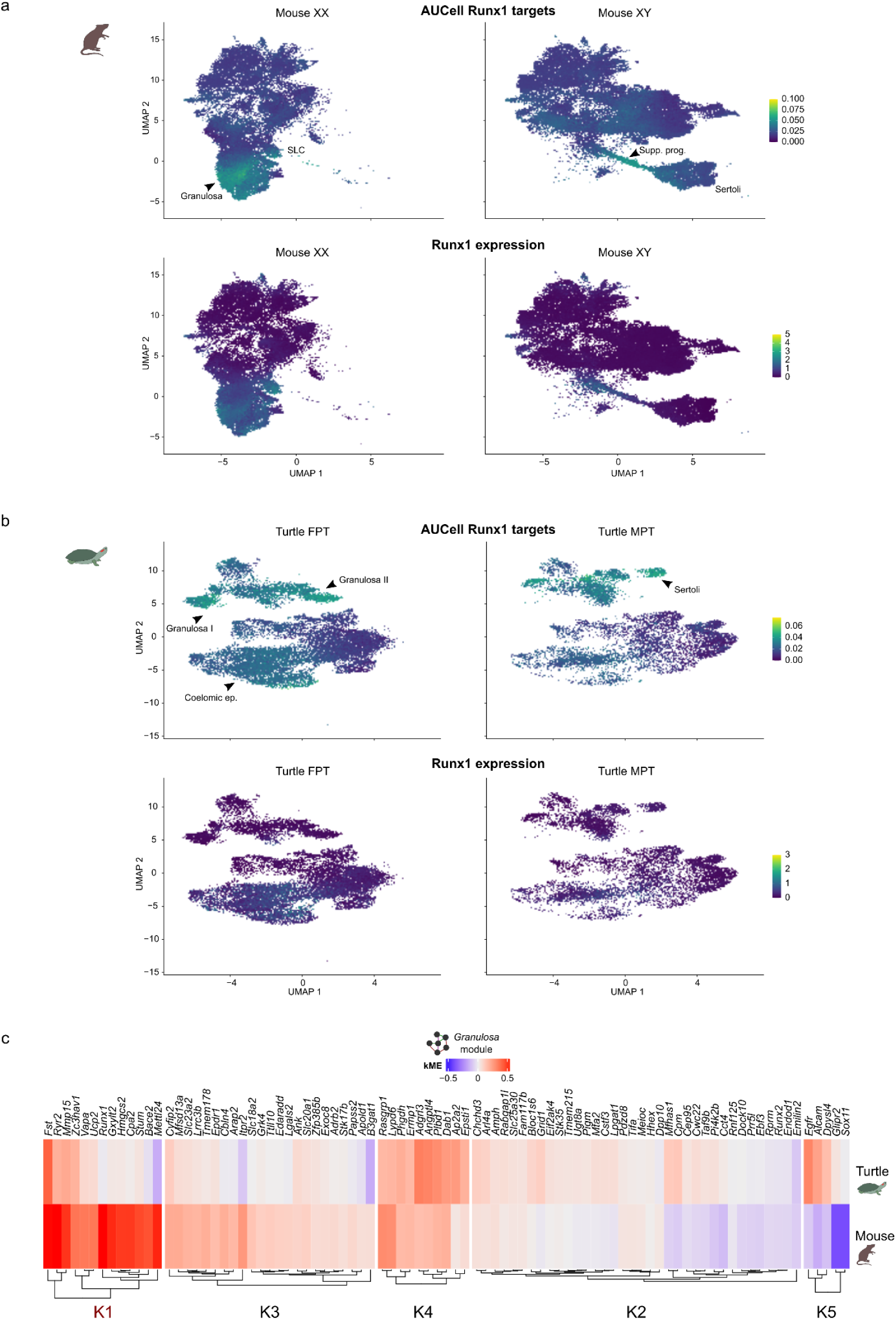
Dismantling of the *Runx1* module in *T. scripta.* a) UMAP reductions of mouse single-cell data with AUCell scores of Runx1 targets in XX (top left) and XY (top right) gonads. Expression of the TF *Runx1* in XX (bottom left) and XY (bottom right) gonadas. Relevant cell types are indicated with arrowheads. b) As in a) but in single-cell from gonads of the turtle *T. scripta*. c) Clustering (k-means) of Runx1 targets according to their membership (kME) in the *Granulosa* module of turtle (top) or mouse (bottom). K1 cluster show hub genes for *Granulosa* in mice that are disconnected in turtles.

**Extended Data Figure 16:**
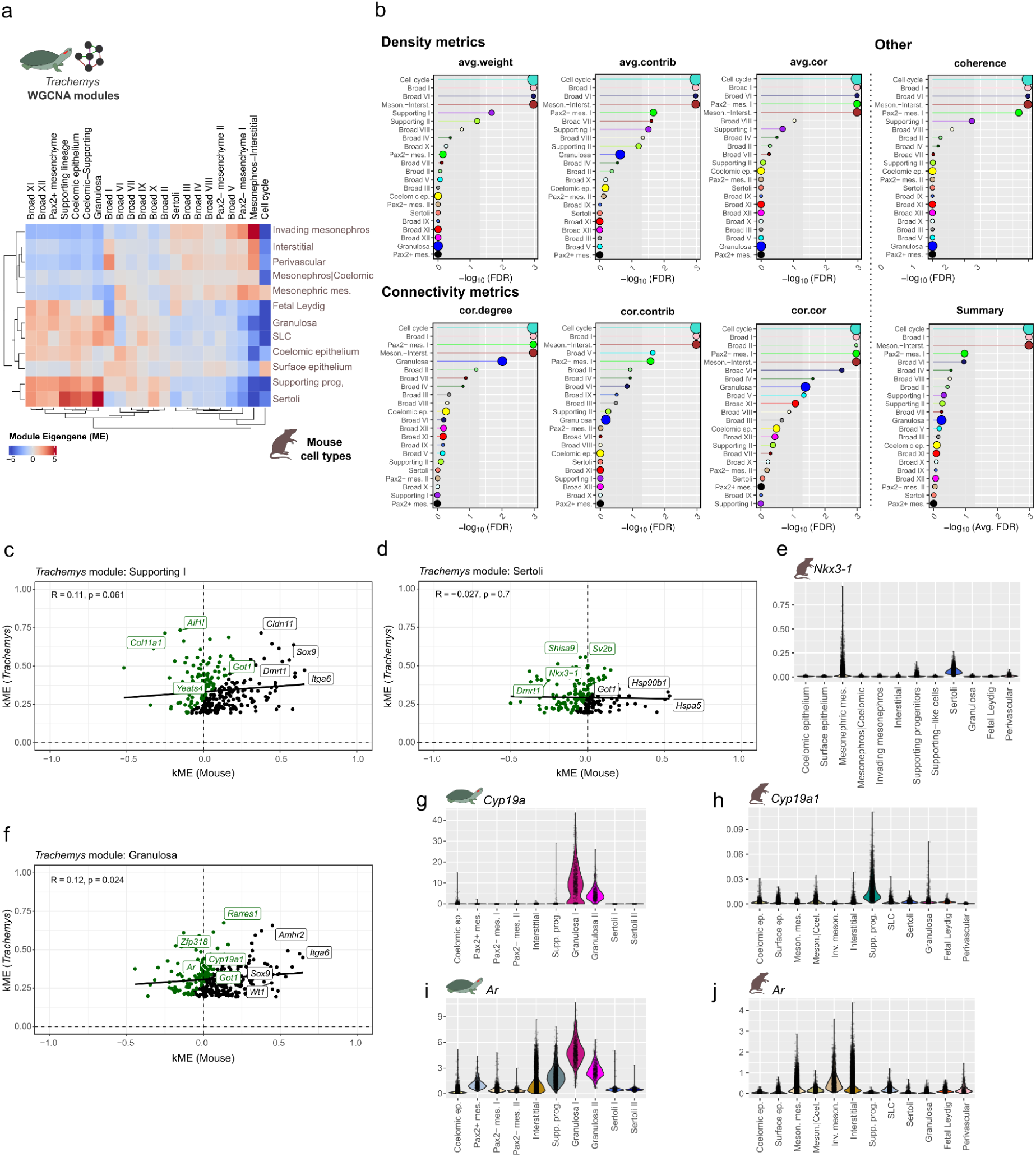
Conservation of *T. scripta* WGCNA modules in mouse. a) Heatmap showing Module Eigengene (ME) scores of *T. scripta* WGCNA modules in each of the cell types of mouse. High ME scores imply that the genes of the *T. scripta* WGCNA module are active in the corresponding cell type of mouse. b) Lollipop plots displaying network conservation FDR values between *T. scripta* and mouse of the different WGCNA modules calculated with NetRep. On the x-axis is represented the negative logarithm of the FDR of a permutation test assessing whether the module has a higher conservation metric than modules generated by chance. See Extended Data Fig. 10. c) Scatterplot showing the membership (kME) of the genes belonging to the WGNCA module *Supporting I* calculated in *T. scripta* both in *T. scripta* (y-axis) and mouse (x-axis). Genes that are associated in *T. scripta* but not in chicken are labeled green (i.e. *Nkx3.1*). d) As in c) but for the *T. scripta* WGCNA module *Sertoli*. e) SCVI normalized expression of *Nkx3-1* in mouse gonadal cell types. f) As in c) but for the *T. scripta* module *Granulosa*. g-h) SCVI normalized expression of *Cyp19a1* in *T. scripta* and mouse gonadal cell types. i-j) SCVI normalized expression of *Ar* in *T. scripta* and mouse gonadal cell types.

**Extended Data Figure 17:**
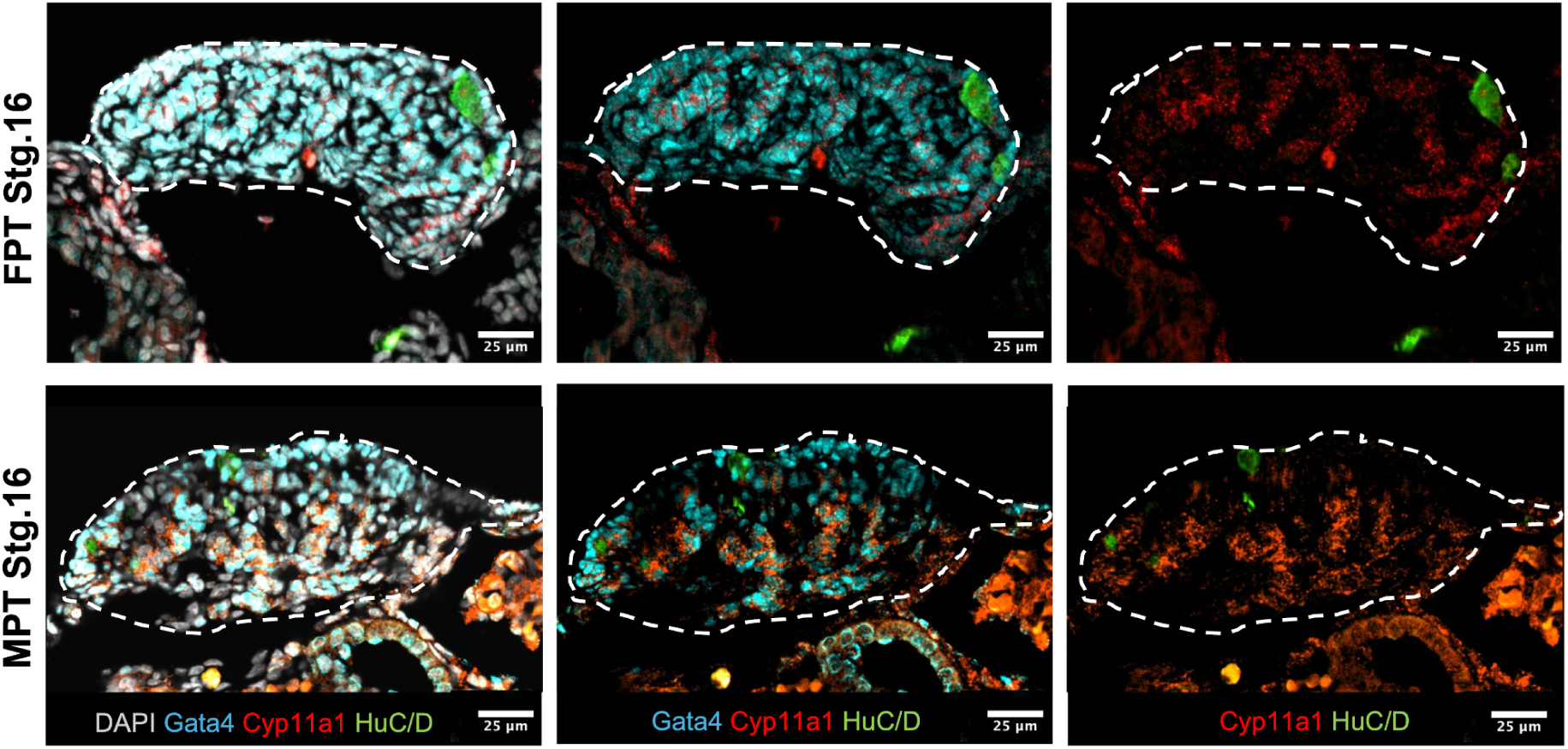
Steroidogenesis in T. scripta gonads. Representative immunofluorescence images of MPT and FPT gonadal cross sections at St.16. DAPI labels the nuclei in grey, HuC/D labels germ cells in green, Gata4 labels gonadal somatic cells in cyan, and Cyp11a1 labels steroidogenic cells in red. Note that Cyp11a1 labels the cytoplasm of Gata4+ cells located in the primitive cords at both temperatures (scalebar, 25 μm). White dashed line outlines the gonad.

**Extended Data Figure 18:**
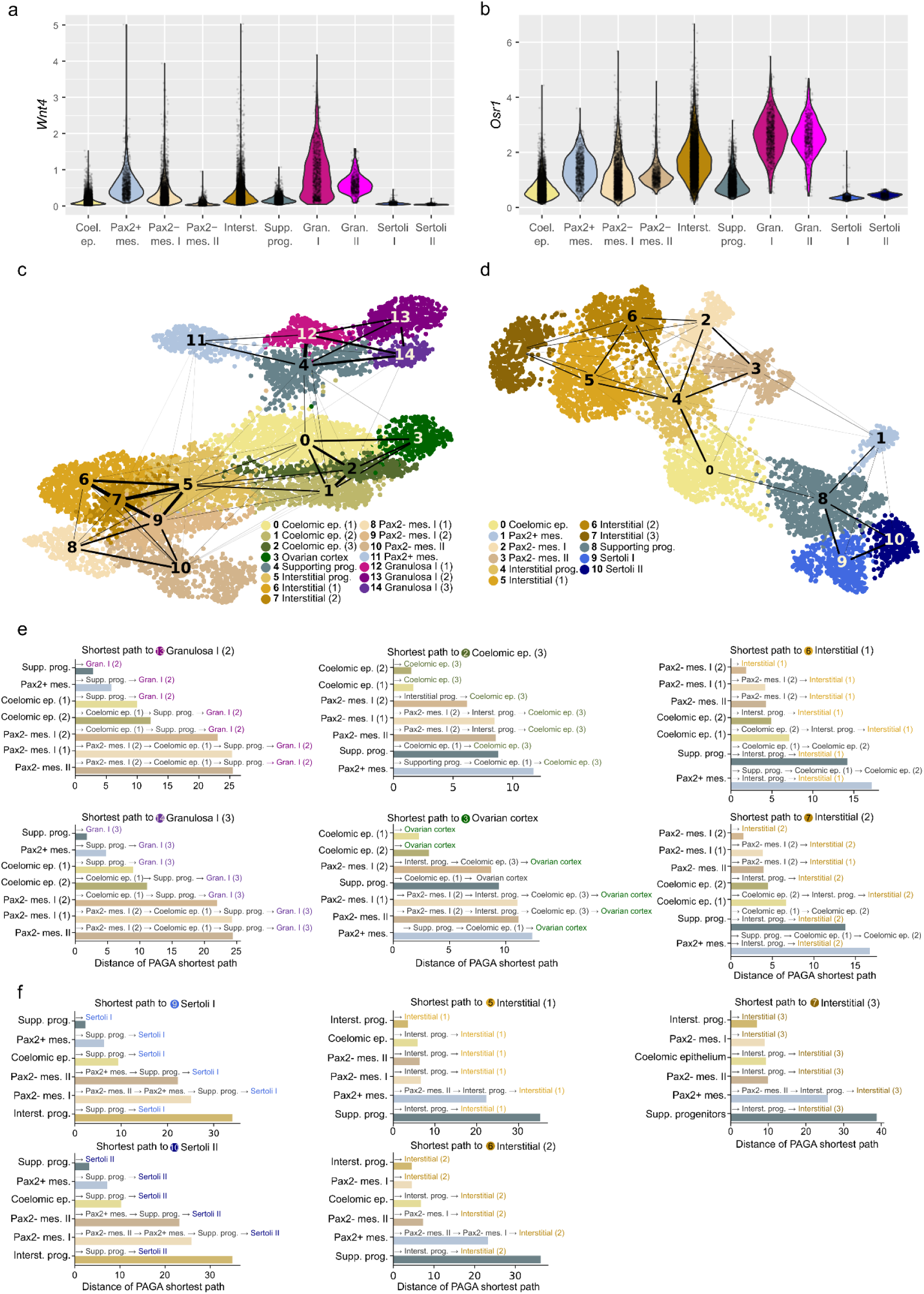
Trajectory analysis in *T. scripta* gonads. a-b) SCVI normalized expression of *Wnt4* and *Osr1*. c-d) UMAP embedding of FPT and MPT somatic gonads respectively, colored according to Leiden subclusters. The width of the connections between clusters represent the PAGA scores. Higher PAGA scores suggest a more likely transition between two cell types. e-f) Barplots represent the shortest distance in the PAGA graph between cell types present in early somatic gonads and terminal cell fates at either FPT or MPT respectively. Shortest distances imply a more likely transition.

**Extended Data Fig. 19:**
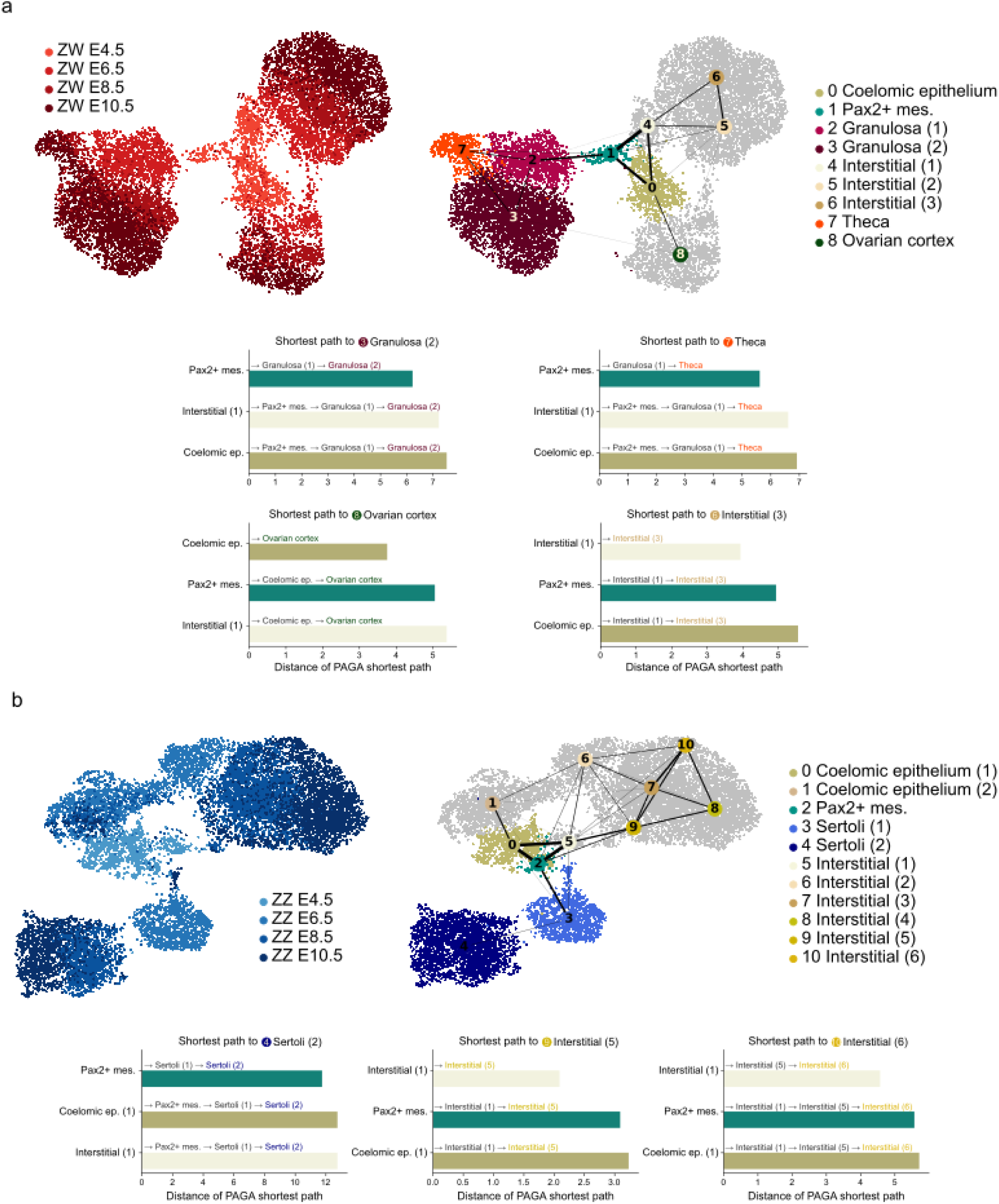
Developmental origin of Sertoli and granulosa cells in chicken. a) PAGA trajectory analysis of female (ZW) somatic gonad cells. At the top left, UMAP embedding from a subcluster of female somatic cells displaying information of the stage at which they were sampled. At the top right, the same UMAP embedding but this time colored according to the Leiden subclusters. The width of the connections between clusters represent the PAGA scores. Higher PAGA scores suggest a more likely transition between two cell types. At the bottom, the barplots represent the shortest distance between early and terminal clusters. Shortest distances imply a more likely transition. b) As in b) but for male (ZZ) somatic gonad cells.

**Extended Data Fig. 20:**
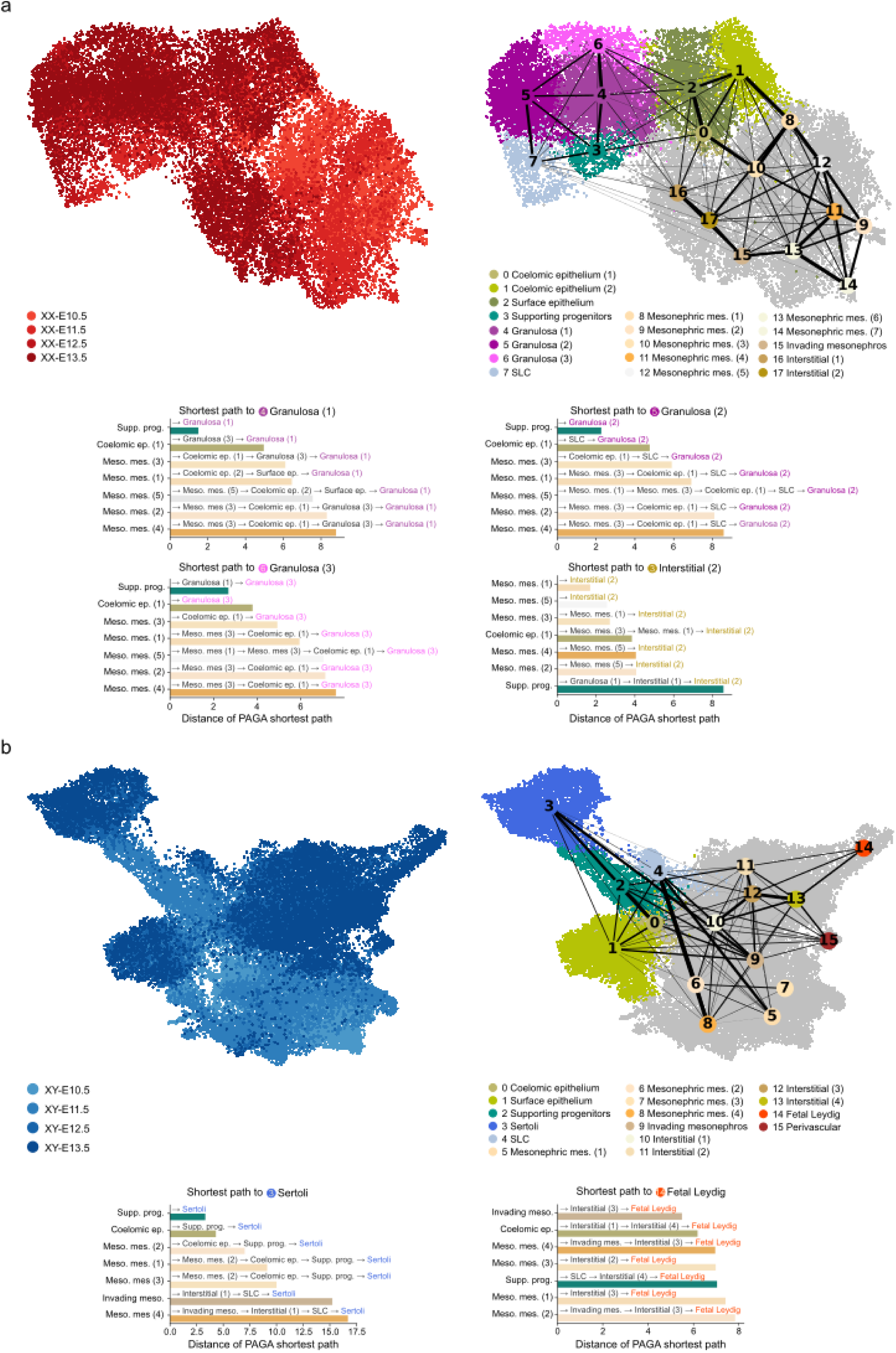
Developmental origin of Sertoli and granulosa cells in mice. a) PAGA trajectory analysis of female (XX) somatic gonad cells. At the top left, UMAP embedding from a subcluster of female somatic cells displaying information of the stage at which they were sampled. At the top right, the same UMAP embedding but this time colored according to the Leiden subclusters. The width of the connections between clusters represent the PAGA scores. Higher PAGA scores suggest a more likely transition between two cell types. At the bottom, the barplots represent the shortest distance between early and terminal clusters. Shortest distances imply a more likely transition. b) As in b) but for male (XY) somatic gonad cells.

**Extended Data Fig. 21:**
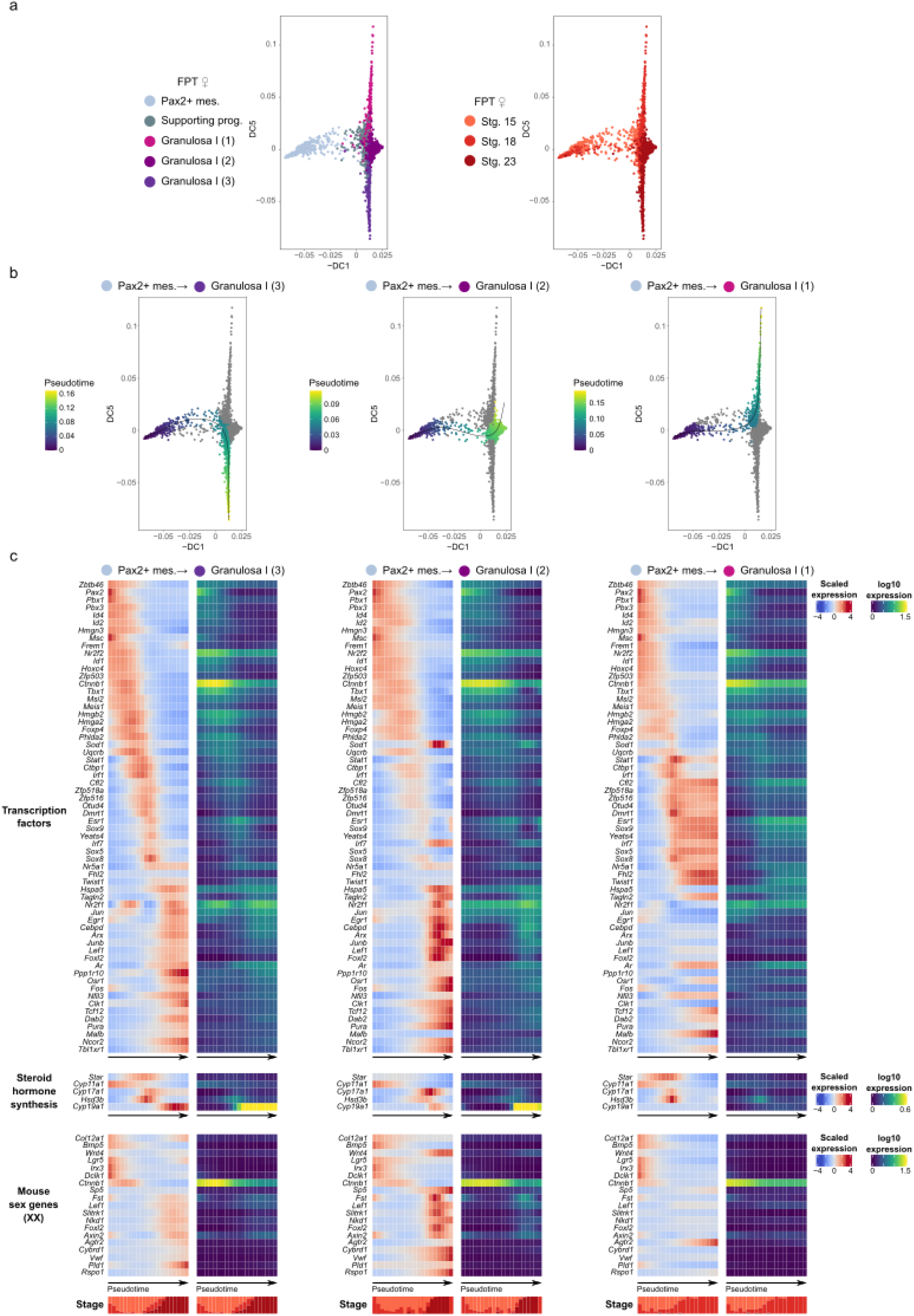
Dynamic expression along the Pax2+ mes. to granulosa differentiation trajectory. a) Diffusion component analysis of cells of the selected clusters in the Pax2+ mesenchyme to granulosa differentiation trajectory. Cells are colored either by cluster (left) or stage (right). b) Same diffusion component reduction as in a) but now cells are colored according to the pseudotime in the differentiation trajectory to three different subclusters of Granulosa I cells. c) Heatmaps showing the scaled (left) and log10 normalized (right) expression of relevant genes along the different pseudotimes. The top heatmaps show the 50 top expressed transcription factors found to be dynamic in either of the three trajectories. The middle heatmaps focus on the expression of steroid hormone enzymes. The bottom heatmaps show a curated set of genes shown to be important for granulosa cell differentiation in mice (Bouma et al. 2010). At the bottom, a stacked barplot shows the proportion of cells of the different stages that are present in each pseudotime window. Darker colors represent later stages.

**Extended Data Fig. 22:**
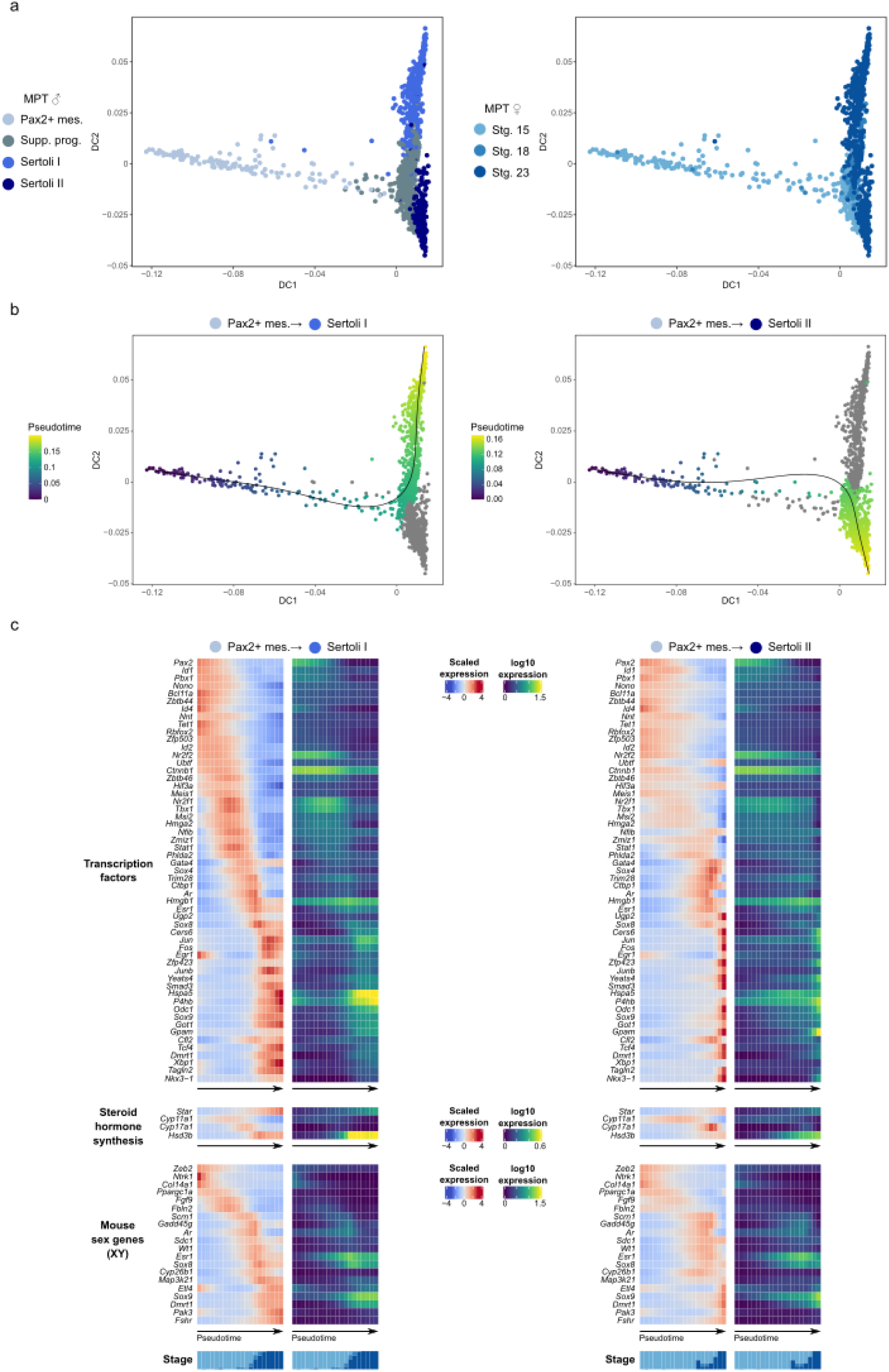
Dynamic expression along the Pax2+ to Sertoli cell differentiation trajectory. a) Diffusion component analysis of cells of the selected clusters in the Pax2+ mesenchyme to Sertoli cell differentiation trajectory. Cells are colored either by cluster (left) or stage (right). b) Same diffusion component reduction as in a) but now cells are colored according to the pseudotime in the differentiation trajectory to either Sertoli I or Sertoli II cells. c) Heatmaps showing the scaled (left) and log10 normalized (right) expression of relevant genes along the different pseudotimes. The top heatmaps show the 50 top expressed transcription factors found to be dynamic in either of the two trajectories. The middle heatmaps focus on the expression of steroid hormone enzymes. The bottom heatmaps show a curated set of genes shown to be important for Sertoli cell differentiation in mice (Bouma et al. 2010). At the bottom, a stacked barplot shows the proportion of cells of the different stages that are present in each pseudotime window. Darker colors represent later stages.

